# A phylotranscriptome study using silica gel-dried leaf tissues produces an updated robust phylogeny of Ranunculaceae

**DOI:** 10.1101/2021.07.29.454256

**Authors:** Jian He, Rudan Lyu, Yike Luo, Jiamin Xiao, Lei Xie, Jun Wen, Wenhe Li, Linying Pei, Jin Cheng

## Abstract

The utility of transcriptome data in plant phylogenetics has gained popularity in recent years. However, because RNA degrades much more easily than DNA, the logistics of obtaining fresh tissues has become a major limiting factor for widely applying this method. Here, we used Ranunculaceae to test whether silica-dried plant tissues could be used for RNA extraction and subsequent phylogenomic studies. We sequenced 27 transcriptomes, 21 from silica gel-dried (SD-samples) and six from liquid nitrogen-preserved (LN-samples) leaf tissues, and downloaded 27 additional transcriptomes from GenBank. Our results showed that although the LN-samples produced slightly better reads than the SD-samples, there were no significant differences in RNA quality and quantity, assembled contig lengths and numbers, and BUSCO comparisons between two treatments. Using this data, we conducted phylogenomic analyses, including concatenated- and coalescent-based phylogenetic reconstruction, molecular dating, coalescent simulation, phylogenetic network estimation, and whole genome duplication (WGD) inference. The resulting phylogeny was consistent with previous studies with higher resolution and statistical support. The 11 core Ranunculaceae tribes grouped into two chromosome type clades (T- and R-types), with high support. Discordance among gene trees is likely due to hybridization and introgression, ancient genetic polymorphism and incomplete lineage sorting. Our results strongly support one ancient hybridization event within the R-type clade and three WGD events in Ranunculales. Evolution of the three Ranunculaceae chromosome types is likely not directly related to WGD events. By clearly resolving the Ranunculaceae phylogeny, we demonstrated that SD-samples can be used for RNA-seq and phylotranscriptomic studies of angiosperms.

## 1 INTRODUCTION

With the recent advances in high-throughput sequencing and analytical methods, phylogenetic reconstruction using genome-wide sequence data has become widely used in plant evolutionary studies (Johnson et al., 2012; Yu et al., 2018). However, whole-genome sequencing of densely sampled phylogenetic analyses has remained impractical and unnecessary due to high costs and computational limitations. Hence researchers have developed reduced-representation methods (e.g., genome skimming, restriction site-associated DNA sequencing or RAD-seq, target enrichment sequencing such as Hyb-seq, and transcriptome sequencing or RNA-seq) as practical tools for phylogenetic studies (Zimmer & Wen, 2015; McKain et al., 2018).

Genome skimming is one of the most widely applied partitioning strategies for phylogenetic inferences, and especially efficient for obtaining complete plastid genome sequences of plants (Dodsworth, 2015; Liu et al., 2018; Zhai et al., 2019; He et al, 2019; Liu et al., 2020; Wang et al, 2020). However, it is often of limited utility to obtain enough single copy nuclear genes for phylogenetic analyses, especially for non-model plant taxa possessing both huge genome sizes and no whole-genome reference (McKain et al., 2018; but see Liu et al., 2021). Hyb-seq is another widely used genome partitioning method that can target low-copy nuclear genes. Like genome skimming, Hyb-seq can use almost all kinds of tissue samples (e.g., silica gel-preserved, flash- frozen, fresh, or even old herbarium materials) (Yu et al., 2018; McKain et al., 2018; Reichelt et al., 2021; Wang et al., 2021). However, Hyb-seq requires a complex laboratory protocol involving bait capture. Furthermore, this method often results in a high proportion of missing data, and also cannot be used to detect ancient whole genome duplication (WGD) based on paralogous genes.

In recent years, using RNA-seq to reconstruct phylogenetic relationships (phylotranscriptomics) and gene family evolution has gained popularity because of its relatively low cost and improved analytical pipelines (Wen et al., 2013, 2015; Wickett et al., 2014; Landis et al., 2017; Zeng et al., 2017; One Thousand Plant Transcriptomes Initiative, 2019; Cheon et al., 2020; Alejo-Jacuinde et al., 2020). Using that method, researchers can often assemble thousands of genes (especially single-copy nuclear genes) from plant taxa under study and use them for both inferring phylogenetic relationships and gene family evolution (Yang and Smith, 2014; Yang et al., 2015; Xiang et al., 2017). Compared to Hyb-seq, RNA-seq uses relatively simple experimental protocols to generate more complete nuclear data, and paralogous genes from RNA-seq data can also be used to infer WGD (McKain et al., 2018).

While improvements in sequencing and extraction protocols have made RNA-seq much easier in plants (Romero et al., 2014; Yang et al., 2017), its application in phylogenetic study still remains challenging. Because RNA is more unstable than DNA, RNA-seq requires more stringent material preservation techniques. Previous studies have used fresh or liquid-nitrogen flash frozen plant tissues or fresh tissue quickly soaked in RNA stabilization solution, and then subsequently preserved in an ultra-low temperature (-80 ℃) freezer (Yu et al., 2018; McKain et al., 2018; Dodsworth et al., 2019). However, because phylotranscriptomic studies often focus on non-model plant taxa, they usually require extensive field work and broad taxon sampling schemes. The logistics of using liquid nitrogen tanks in the field or expensive RNA stabilization solution to preserve collected plant tissues at multiple locations, not to mention quick access to an ultra-low temperature freezer for subsequent laboratory work, greatly limits the practicality of using RNA-seq for phylogenetic studies (Zimmer & Wen, 2015; Yang et al., 2017).

Traditional DNA-based molecular phylogenetics, DNA barcoding, as well as genome skimming and RAD-seq methods, often use silica gel-dried leaf tissues (Narzary et al., 2015; Yu et al., 2018). Such sampling method is cheaper and amenable to collecting and transporting large number of samples. Much emphasis has been placed on how different plant tissues, preservation methods, and RNA extraction protocols may impact the quantity and quality of extracted RNA (Johnson et al., 2012; Romero et al., 2014; Yang et al., 2017), but no empirical study has explored silica gel-dried plant tissues for RNA-seq. This may be due to the reasons that even though total RNA may be extracted from silica gel-dried plant tissues, it is not possible to quantitatively measure gene expression using this kind of samples for evo-devo studies. However, a phylotranscriptomic study needs to obtain large nuclear data sets for pertinent plant taxa for analysis, and it does not need to quantitatively measure gene expression. Therefore, if silica gel-dried and moderately frozen (-20 ℃) plant tissues can be sequenced via RNA-seq, it may greatly expand the utility of RNA-seq in phylogenetic studies.

As a member of the early-diverging eudicots, the family Ranunculaceae consists of more than 50 genera and 2,000 species, and it has been widely used as a model for angiosperm evolutionary studies (Tamura, 1995; Emadzade et al., 2010; Johansson and Jansen 1993; Ro et al. 1997; Wang et al., 2016; Zhai et al., 2019). Systematists have also long focused on the family (Wiegand 1895; Smith 1926; Langlet 1927, 1932; Wodehouse 1936; Gregory 1941; Tamura, 1995; Ro et al., 1997; Zhai et al., 2019), and it has been used as a model for evolutionary studies addressing questions on floral development, genome duplication, and key innovations (Kramer et al., 2003, Kramer, 2009; Rasmussen et al., 2009; Kramer and Hodges 2010; Galimba et al., 2012; Zhang et al., 2013; Wang et al., 2015; Damerval and Becker, 2017; Aköz and Nordborg 2019; Shi and Chen 2020).

Traditionally, systematic studies of the Ranunculaceae emphasized morphological characters (especially floral traits and fruit types) (Prantl, 1887). Because of its complex evolutionary history, the infrafamilial classifications of Ranunculaceae have remained highly controversial (Prantl, 1887; Hutchinson, 1923; Takhtajan, 1980; Cronquist, 1981; Tamura, 1995). Since the studies by Langlet (1927, 1932), cytological characters, including chromosome size and base number, have become important for delimiting subfamilies and tribes of the family (Janchen, 1949; Tamura, 1995). Since the 1990s, many molecular phylogenetic studies of the Ranunculaceae based on Sanger sequencing (Johansson and Jansen, 1993; Hoot, 1995; Johansson, 1995; Ro et al., 1997; Hoot et al., 2008; Wang et al., 2009, 2016; Cossard et al., 2016) have helped clarify many systematic issues, including the delimitation of the family and its phylogenetic framework. Yet detailed tribal relationships within the family have remained controversial and conflicts among different gene trees were common in the family. Recently, phylogenomic studies using complete plastid genome data have greatly improved the resolution and statistical support of many previously unresolved clades within the family (Zhai et al., 2019; He et al., 2019). However, because plastid genome sequences are uniparentally inherited, they may not truly reflect phylogenetic relationships among taxa (Greiner et al., 2015). Therefore, the tribal relationships, especially within the core Ranunculaceae (defined below), still need further clarifications using multiple single-copy nuclear markers.

Using molecular data, Wang et al. (2009, plastid and nrDNA sequences) and Zhai et al. (2019, complete plastid genome sequences) resolved 14 tribes within the Ranunculaceae. Except for three early diverged tribes (Glaucideae, Hydrastideae, and Coptideae) that have *Coptis* type (C-type) chromosomes, the remaining 11 tribes formed the core Ranunculaceae clade. Within the core Ranunculaceae clade, there are two major chromosome types: the *Ranunculus* type bearing large chromosomes with a base number of 8 (R-chromosome group, or R-type) and the *Thalictrum* type having short, small chromosomes with a base number of 7 or 9 (T-chromosome group, T-type) (Gregory, 1941). The genome sizes between these two chromosome types vary 180- fold, with the smallest genome size (1C value: 240.10 MB) in *Thalictrum rhychocarpum* (T-type chromosome) and the largest genome size in *Hepatica nobilis* (R-type chromosome, 1C value: 43,708 MB) (https://cvalues.science.kew.org/). The C-type chromosome genome sizes, from 1,150 MB (*Coptis*, Liu et al., 2021) to 1,310 MB (*Hydrastis*, https://cvalues.science.kew.org/), lie between those of the T- and R-type chromosome taxa, but closer to the T-type.

Genome size variations across eukaryotic taxa often result from macro-mutations, such as WGDs and repetitive element proliferation (Marburger et al., 2018), and WGD provides raw materials for evolutionary novelty in seed plants and has been widely reported in angiosperms (Ren et al., 2018; Cai et al., 2019; One Thousand Plant Transcriptomes Initiative, 2019; Park et al., 2020). Although several recent studies have discussed WGD in some Ranunculaceae taxa (Park et al., 2020; Xie et al., 2020; Liu et al., 2021), their sampling was limited. Until now, the lack of a well-sampled phylogenetic framework of Ranunculaceae impedes our ability to understand the evolutionary processes of key characters, such as floral and fruit characters. Moreover, the exploration of genome-wide gene tree discordance is essential for understanding the underlying evolutionary processes of Ranunculaceae. We also need a robust phylogeny within which to investigate how the three chromosome types evolved and whether ancient WGD events were important in that evolution.

Here, we extracted RNA from both liquid-nitrogen frozen and silica gel-dried leaf tissues and conducted a phylotranscriptomic study of Ranunculaceae. We compared the quality and quantity of extracted RNA between liquid-nitrogen and silica-preserved samples, assessed whether silica-dried plant tissues can be used for phylogenomic study, used multiple single-copy nuclear datasets to reconstruct the Ranunculaceae phylogenetic framework, analyzed the discordance among the gene trees (nuclear as well as cyto-nuclear discordance), and explored divergence times and WGD events in Ranunculaceae and its closely related taxa.

## 2 MATERIALS AND METHODS

### 2.1 Plant Material

Because previous phylotranscriptomic studies mainly used transcriptome data from vegetative tissues (Johnson et al., 2012; Zeng et al., 2017; One Thousand Plant Transcriptomes Initiative, 2019), we likewise collected 27 leaf samples from 22 genera of Ranunculales (20 Ranunculaceaae and two Circaeasteraceae genera) for our study. Among them, 21 leaf samples were dried with silica gel (SD-sample) and the other six were liquid nitrogen-preserved leaf tissues (LN-sample). According to other studies (Johnson et al., 2012; McKain et al., 2018), young developing leaf tissues of the 21 SD- samples were dried in blue Tel-Tale silica gel (Sinopharm Chemical Reagent Co. Ltd., Shanghai, size of particles: 2-8 mm), which are widely used in preserving DNA for plant molecular phylogenetic studies. Once completely dried, the leaf tissues were deposited in a -20 ℃ freezer. The six LN-samples (also young developing leaf tissues) were quickly deposited in liquid nitrogen and then ground for RNA extraction. Voucher specimens were deposited in the Herbarium of Beijing Forestry University (Table 1).

**Table 1.**
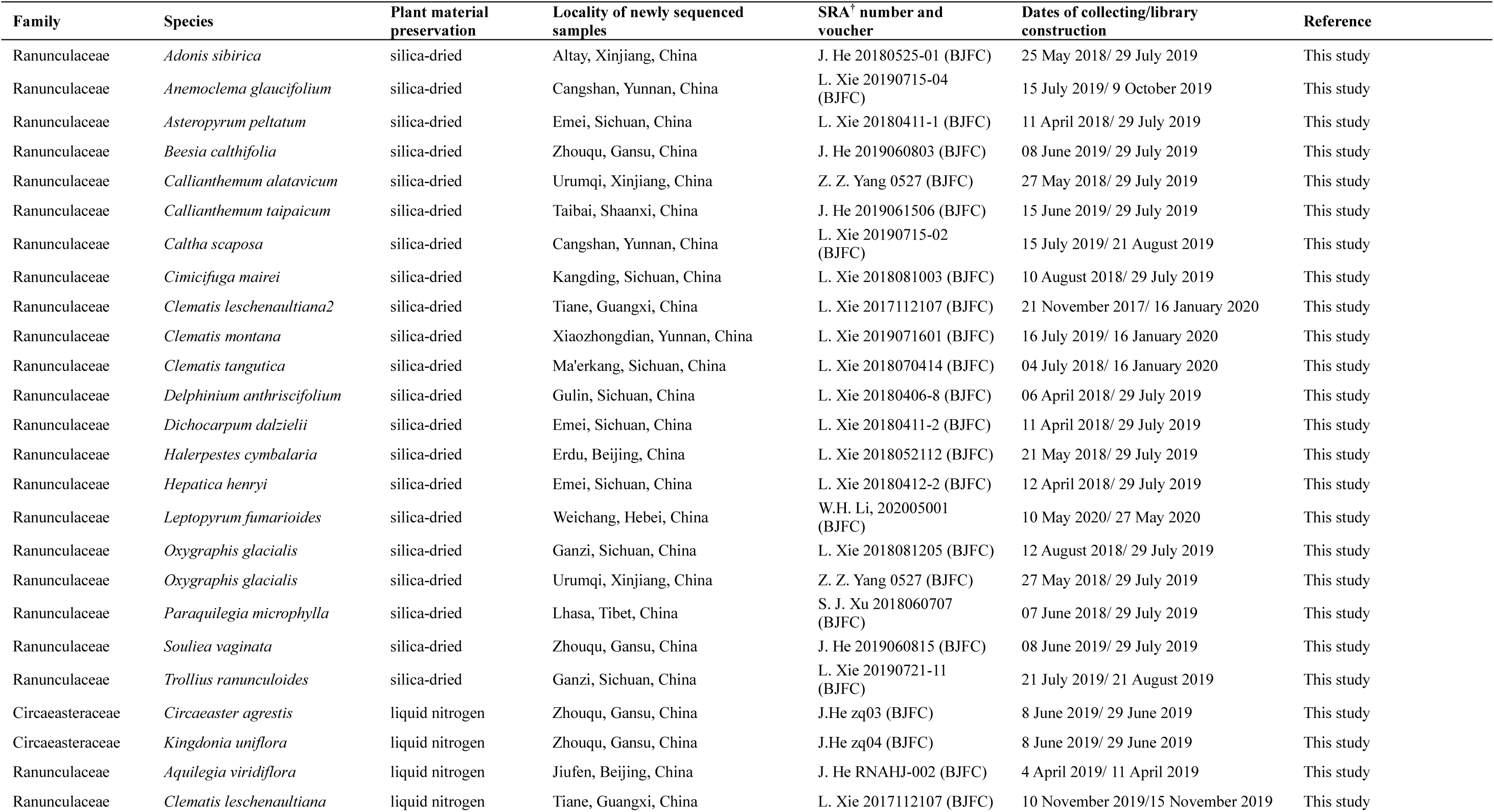
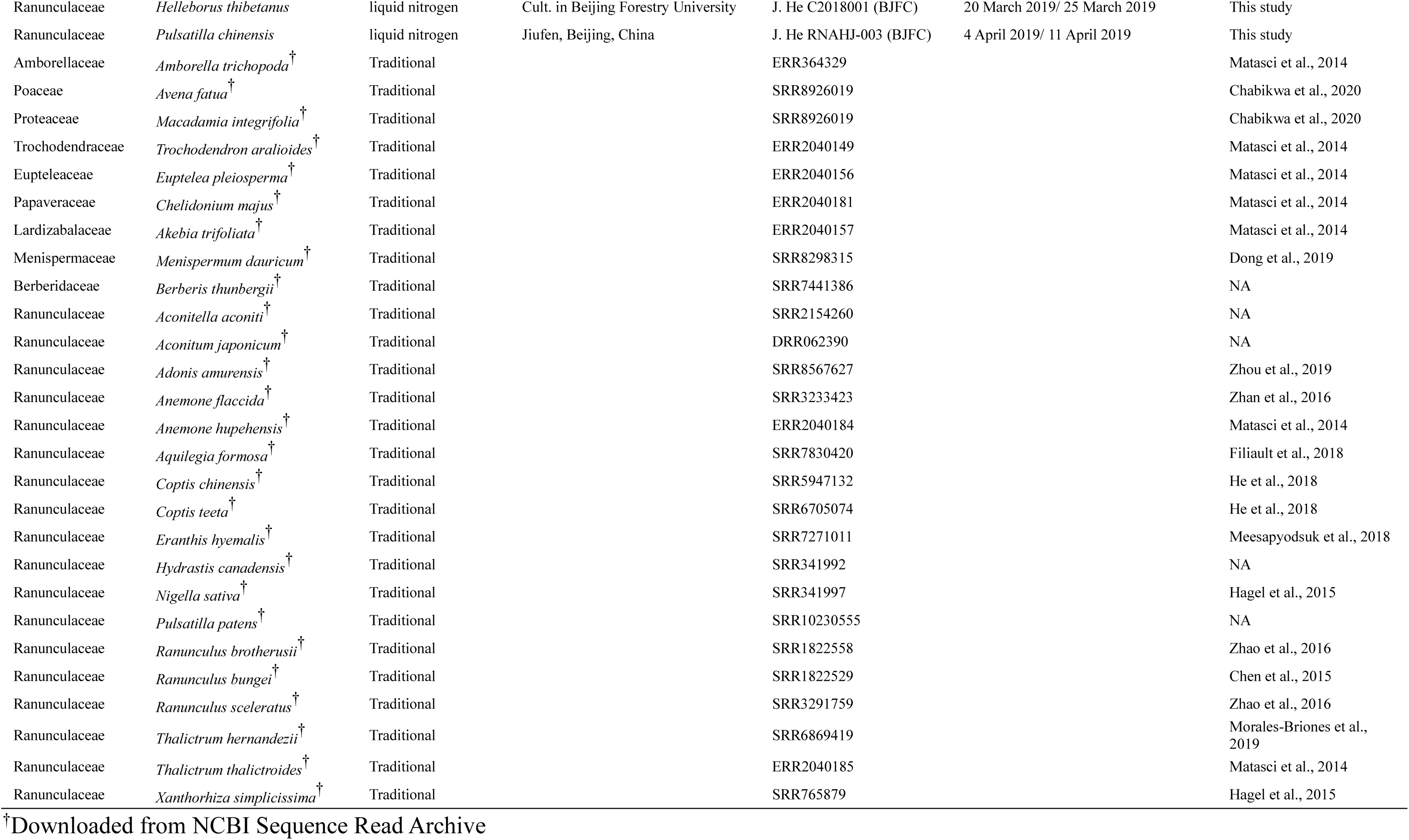
Information about samples used in this study

Next, we downloaded 27 published transcriptomes from GenBank—18 from the Ranunculaceae, six from other Ranunculales families, and three from distantly related angiosperms that served as outgroups. Our sampling covered 13 of all 14 Ranunculaceae tribes (Wang et al., 2009, 2016; Zhai et al., 2019), only the first diverged clade, Glaucideae, was not available for the study. All 27 downloaded transcriptomes were obtained from either freshly collected, liquid-nitrogen-preserved, or RNA stabilization solution-treated leaf tissues. We refer to them as the traditionally treated samples, T-samples.

### 2.2 RNA extraction and sequencing, and cleaning raw reads

We sent our collected leaf tissues to Biomarker Technologies Corporation (http://www.biomarker.com.cn) for RNA extraction, library construction, and next generation sequencing as follows. For each SD- and LN-sample, 100 mg leaf tissues were ground and then total RNA was extracted using a conventional TRIzol-based RNA extraction kit (TRIzon, CoWin Biosciences, Jiangsu, PR China) following the manufacturer’s protocol (see Suplementary Materials). Then, the extracted RNA samples were run on standard RNase-free 1.0% agarose gels and measured using a NanoDrop 2000 spectrophotometer (ThermoFisher Scientific, Waltham, Massachusetts, USA) at 230, 260, and 280 nm to assess the purity and intactness of the samples. Subsequently, the OD260/280 and OD260/230 ratios were calculated. RNA integrity numbers (RIN) and rRNA ratios (28S/18S) were detected by a 2100 Bioanalyzer (Agilent Technologies, Santa Clara, California, USA). Next, the cDNA library was generated from the total RNA extracts by using reverse transcription PCR. The libraries were sequenced on a NovaSeq 6000 platform (Illumina, San Diego, California, USA), ultimately yielding about 6 Gb of raw data per sample. To obtain high-quality clean data, the FASTX-Toolkit (http://hannonlab.cshl.edu/fastx_toolkit/download.html) was used to remove adaptors and low-quality reads from the raw reads. All raw reads were deposited in the NCBI Sequence Read Archive (SRA, BioProject: PRJNA739247).

The 27 downloaded transcriptomes retrieved from the NCBI SRA database were unzipped using the sratoolkit v.2.9.2 (https://ncbi.github.io/sra-tools/). Low-quality bases were removed using Trimmomatic v.0.38 (Bolger et al., 2014).

### 2.3 Transcriptome assembly, gene family clustering, and inferring orthology

The 54 clean transcriptome data were *de novo* assembled using Trinity v.2.5.1 (Grabherr et al., 2011) with default parameters, and the longest isoforms of related contigs were selected using the Trinity helper script (get_longest_isoform_seq_per_trinity_gene.pl). To reduce redundancy, we ran the raw assemblies through CD-HIT v.4.6.2 (Fu et al., 2012) with parameters -c 0.95 and -n 8 (Feng et al., 2019) and then assessed each assembly’s completeness by comparing them against single-copy orthologs in the BUSCO database (Simão et al., 2015). Coding regions of each putative transcript were predicted using TransDecoder v.5.0 (https://github.com/TransDecoder/TransDecoder/releases).

To assign the *de novo* assembled sequences into single copy orthologous gene (SCOG) families, we used an integrated method proposed by Cai et al. (2019) that rigorously takes sequence similarity and species phylogeny into account. To begin, we first constructed transcriptome homology scans using Proteinortho v.6.0.10 (Lechner et al., 2011) in the Diamond mode (Buchfink et al., 2015) and then searched the resulting clusters to identify gene families containing at least 38 in-group species (> 70%), which then obtained 3,634 candidate homologous clusters.

To reduce errors that sometimes occur in this similarity-based homology search because of deep paralogs, mis-assembly, or frame shifts (Yang & Smith, 2014), we ran multiple scans to increase statistical and inferential powers. Since the SCOGs could have been clustered not only from nuclear genomes but also from organellar genomes, we deleted all the organellar SCOGs by using a BLAST analysis against a *Clematis* chloroplast genome (NC039579) and a *Nelumbo* mitochondrial genome (NC030753).

Considering that a sequence clustered to an erroneous ortholog often shows an unexpectedly long branch in the gene tree (Mai and Mirarab, 2018), we used Treeshrink v.1.3.9 (Mai and Mirarab, 2018) to scrutinize each gene tree, filtering out sequences with long branches, and deleting them from the alignments.

Each gene tree was generated by Treeshrink v.1.3.9 in the following four steps: (1) align the peptide sequences of each putative SCOG with MAFFT v.7.475 (Katoh and Standley, 2013); (2) convert the alignments into the corresponding nucleotide alignments; (3) delete each nucleotide site with more than 20% missing data by using a python script (https://github.com/Jhe1004/DelMissingSite); and (4) reconstruct gene trees using a maximum likelihood (ML) method implemented in RAxML v.8.2.12 (Stamatakis, 2014) and the GTR+G model with 100 bootstrap replicates. Sequences that generated long branches in gene trees were removed from the alignments (Mai and Mirarab, 2018) and then, using a python script (https://github.com/Jhe1004/ConcatenateAlignment), the remaining alignments were concatenated into a super-alignment for further analysis.

Finally, to improve the accuracy of gene trees, we reconstructed the Ranunculaceae phylogenetic framework using three different strategies. (1) Limit the minimum length of each SCOG alignment (Yang et al., 2015). (2) Limit each SCOG tree’s minimum average bootstrap values (Molloy and Warnow, 2018). (3) Limit the allowed maximum of missing taxa for each SCOG alignment (Yang et al., 2015). For each strategy, we set two different screening thresholds (described below).

We obtained eight data sets for phylogenetic analysis: (1) 3,611 concatenated SCOGs (SCOG-concatenated), (2) 3,611 separate SCOGs (SCOG-coalescent), (3) SCOGs each with at least 1,000 bp (SCOG-1000bp), (4) SCOGs each with at least 2,000 bp (SCOG-2000bp), (5) SCOGs each with at least 50% mean bootstrap values (SCOG-50BS), (6) SCOGs each with at least 80% mean bootstrap values (SCOG-80BS), (7) SCOGs each with no more than 20% missing taxa (SCOG-20MS), and (8) SCOGs each with no more than 10% missing taxa (SCOG-10MS).

### 2.4 Phylogenetic reconstruction

For phylogenomic analysis, we used both coalescence- and concatenation-based methods for each data set. For the coalescent-based method, the Accurate Species Tree Algorithm v.4.4.4 (ASTRAL; Zhang et al., 2018) was applied. Single-gene ML trees for seven of the data sets (not the SCOG-concatenated data set) were reconstructed in RAxML v.8.2.12 on the CIPRES Science Gateway v.3.3 (Miller et al., 2010) using the GTR+G model. All those trees were used as input in ASTRAL for species tree inference. For the concatenation method, all the SCOG-concatenated data sets were calculated for using ML method implemented in RAxML v.8.2.12 with the GTR+G model and bootstrap percentages computed after 200 replicates. In order to test cyto-nuclear discordance, the plastome sequence data were obtained from previously published studies (Table S1) and the plastome phylogeny was inferred using the ML method as described above.

### 2.5 Molecular dating

The molecular clock estimation used the SCOG-concatenated data set and constrained with the species tree topology. We used three widely accepted fossil calibrations in our dating analysis. (1) The Ranunculales stem group (the tree root) was constrained at a minimum age of 113 Mya (range = 113–125 Mya), which was based on the earliest (early Cretaceous) mega-fossil (flower) record for the order, *Teixeiria lusitanica* (von Balthazar et al., 2005; Magallón et al., 2015). (2) *Prototinomiscium vangerowii* gave the stem age of the Menispermaceae (Knobloch and Mai, 1986; Kadereit et al., 2019) at a minimum of 91 Mya (range = 91–99 Mya). (3) The stem of the *Actaea* clade was constrained at a minimum age of 56 Mya (range = 56–67 Mya), based on the fossil record of *Paleoactea nagelii* (Pigg and DeVore, 2005; Wang et al., 2016).

Divergence times were estimated using MCMCTree implemented in the PAML v.4.9a package (Yang, 2007), and it was run in two steps. First, the topology was constrained by the ASTRAL species tree; then the ML estimates of the branch lengths, the gradient vector, and the Hessian matrix were generated using the codeml model from the PAML v.4.9a package (Yang, 2007). Then, those variables and data (Hessian matrix) were fed into MCMCTree control files, together with the GTR substitution model. An autocorrelated relaxed clock model (Rannala and Yang, 2007) was used to estimate the divergence time posterior distribution and the MCMC chain ran for 100,000,000 generations, with tree sampling every 1,000 generations after a burn-in of 25,000,000 iterations. We used FigTree v.1.4.4 (http://tree.bio.ed.ac.uk/software/figtree) to generate the chronogram.

### 2.6 Visualizing and assessing conflicts among the gene trees

We used Phyparts v.0.0.1 (Smith et al., 2015) to explore conflicts among the nuclear gene trees. To reduce computational time, we mapped gene trees from the SCOG- 1000bp data set onto the species tree generated by the SCOG-coalescent data set, and the numbers of discordant and concordant nodes of the gene trees compared to those of the species tree were then counted and summarized. To further visualize conflicts, we built cloud tree plots using the python package Toytree v.2.0.5 (Eaton, 2020). Because cloud tree plots cannot accommodate missing taxa among gene trees, we used the python package ETE3 v.3.1.2 (Huerta-Cepas et al., 2016) to filter and prune the gene trees to include all Ranunculaceae samples and only one outgroup (*Berberis thunbergii*). Ultrametric gene trees were time-calibrated with TreePL v.1.0 (Smith and O’Meara 2012) and, using the python package DendroPy v.4.5.2 (Sukumaran and Holder 2010), each gene tree’s nodes were calibrated by age estimates extracted from the MCMCTree dating results described above. Ultimately, we obtained 92 ultrametric gene trees for cloud tree plots.

### 2.7 Coalescence simulations

Discordance among gene trees is likely caused by incomplete lineage sorting (ILS), hybridization, or other sources of error and noise (Sang & Zhong, 2000; Roos et al., 2011; Morales-Briones et al., 2021). Among them, ILS has attracted great attention in previous studies (Edwards 2009), and several inference methods have been developed to accommodate ILS as the source of discordance (Edwards et al. 2016). To evaluate if ILS alone could explain the gene tree discordance in this study, we performed a coalescent simulating analysis following published studies (Wang et al., 2018; Yang et al., 2020; Morales-Briones et al., 2021). If the coalescent model fit the empirical gene trees well, the simulated gene trees from that model would be consistent with the empirical gene trees and ILS most likely explained the gene tree incongruence well. We used the function “sim.coaltree.sp” implemented in the R package Phybase v.1.5 (Liu and Yu, 2010) to simulate 10,000 gene trees under the multispecies coalescent (MSC) model. After the input ASTRAL ultrametric species tree was modified using a Perl script (Wang et al., 2018), we calculated the distribution of tree-to-tree distances (Robinson and Foulds, 1981) between the ASTRAL species tree and each empirical gene tree (SCOG-1000bp data set) using the Python package DendroPy v.4.5.2 (Sukumaran and Holder 2010). We then compared the distribution of distances between the ASTRAL species tree and the simulated gene trees. Because that calculation requires both trees to have the same set of taxa, we only kept 13 tribal (*sensu* Wang et al., 2016) representatives of the Ranunculaceae for this analysis.

We also carried out a coalescent simulation (Rose et al., 2021) to evaluate if ILS alone could explain cyto-nuclear discordance in Ranunculaceae. If ILS explained the cyto-nuclear discordance, a number of simulated gene trees would be consistent with the plastid genome tree. So, using the R package Phybase v.1.5, we simulated 10,000 gene trees from our ASTRAL species tree under the MSC model. Then we used Phyparts v.0.0.1 to explore conflicts among the simulated gene trees and the plastid genome tree. The proportion of gene tree concordance compared to the plastid genome tree was then counted and summarized.

### 2.8 Species network analysis

We used a Julia package, PhyloNetworks (Solís-Lemus et al., 2017), that models ILS and gene flow using a maximum pseudolikelihood method (Yu and Nakhleh, 2015) to investigate high probability hybridization events and to calculate γ, the vector that reflects the proportion of genes inherited from each parent in a hybrid lineage. Due to computational restrictions, the package cannot run large numbers of samples, so we kept the 13 tribal representatives of the Ranunculaceae for this analysis. From the input gene subtrees extracted from the SCOG-1000bp data set, we obtained 614 gene subtrees for the 13 taxa and, in PhyloNetworks, performed SNaQ (Species Networks applying Quartets) analyses that allowed for zero to five maximum hybridization events with ten independent runs. The best number of hybridization events was selected by plotting the pseudolikelihood scores. A sharp score increase is expected until it reaches the best number, after which the score increases slowly. Finally, we used 100 bootstrap replicates to perform hybrid branch support on the best network.

### 2.9 Detecting and dating WGD in the Ranunculaceae

In this study, WGD events were detected and located using two complimentary methods: a tree-based method to infer WGD and the absolute dating method to verify that result. For the tree-based method, input gene family trees were constructed from all 54 samples by using Proteinortho v.6.0.10 (Lechner et al., 2011) with a cut-off of E = 1 × 10^−5^. For each gene family, ML trees were estimated by RAxML v.8.2.12 under the GTR+G model and using a customized Python script (https://github.com/Jhe1004/reroot). The RAxML analysis output file (bestTree file) was re-rooted and transformed into an input file for the next step. Tree-based WGD inference was conducted using PhyParts v.0.0.1, which mapped the gene family trees to the species tree (based on the SCOG-coalescent data set using ASTRAL). After that, we recorded the nodes that contained duplication events when the descendants of the node had multiple gene copies for at least two terminal samples (Smith et al., 2015).

For the dating analyses, we followed methods of Vanneste et al. (2014). We used Proteinortho v.6.0.10 (Lechner et al., 2011) to obtain homologous gene families (including both orthologous and paralogous gene families) among the Ranunculaceae samples, ultimately finding 12,332 gene families that contain at least two paralogous copies. Then, those gene families were aligned using MAFFT v.7.475 and ML trees were reconstructed by RAxML v.8.2.12 under the GTR+G model. Next, we used TreePL (Smith & O’Meara, 2012) to date the divergence times of each gene family. For this analysis, we constrained both the root and the crown ages of the clade (Berberidaceae, Ranunculaceae) by using the time estimations found in our MCMCTree dating analysis. Then, we filtered the gene trees to ensure that Ranunculaceae and Berberidaceae were sister groups. After all the gene families had been calculated, divergence time estimates of gene duplicates were extracted and WGD curves were plotted using Gaussian mixture models in the Python package sklearn v.0.24.1 (Pedregosa et al., 2011).

## 3 RESULTS

### 3.1 RNA-seq quality from silica gel -dried and traditionally treated tissues

The mean values of some RNA quality and quantity reference indices of LN- samples were higher than those of SD-samples (e.g., RIN, 28S/18S, and OD260/280), but were lower for other indices (e.g., OD260/230, total RNA, and RNA concentration) (Figure 1 a-b, Table S2). However, one-way analysis of variance (ANOVA) showed that none of those differences was significant at *P* < 0.05.

**Figure 1.**
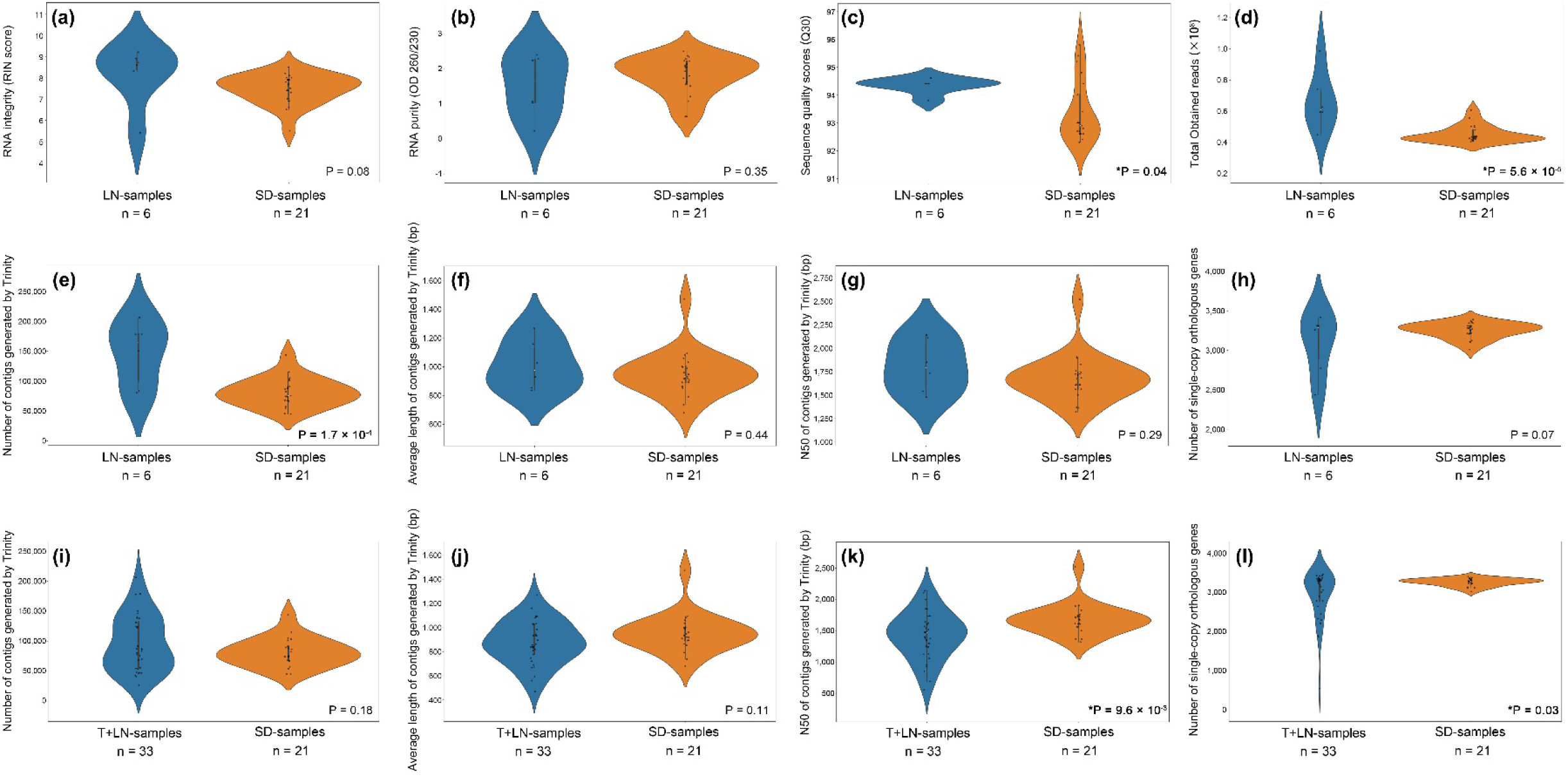
Quality and quantity assessments of RNA extracted from Ranunculales leaf tissues preserved in one of three different ways. Comparisons are of (a-d) genomic RNA, (e-g and i-k) de-novo sequences assembled by Trinity v.2.1.5 (Grabherr et al., 2011), and (h and l the numbers of single-copy orthologous genes compared to BUSCO (Simão et al., 2015). The LN+T samples category includes the 27 samples downloaded from GenBank (see Table 1) plus the liquid nitrogen-preserved samples (n = 6). One-way ANOVA, * *P* < 0.05. OD: optical density; Q30: Phred quality score 30 threshold. For detailed sampling information see Table 1

We then counted and compared the obtained reads, bases, Q20, Q30, and GC content of each treatment. Except for GC content, the other reference indices of LN- samples were significantly better than those of SD-samples (Figure 1 c-d, Table S2). We further assembled the transcripts for all the samples and determined the numbers and average lengths of the contigs (NC and ALC, respectively), the N50s, and the completeness of the assemblies and compared them to BUSCO (Figure 1e-h, Table S3). The results showed that, except for the mean value of NC, which is significantly lower in SD-samples than in LN-samples, all the reference indices were not significantly different between the two treatments.

We next pooled the transcriptome data of our six LN-samples with that of the 27 downloaded T-samples (collectively called T+LN-samples) and compared their combined NC, ALC, and N50 and the completeness of their assemblies, compared to BUSCO, to those indices of the SD-samples (Figure 1i-l, Table S4). The results demonstrated that SD-samples had significantly higher N50 than the T+LN-samples had (Figure 1k, *P* < 0.05, Supplementary Table S4), but there were no statistically significant differences between the NC, ALC, and BUSCO comparison indices (Figure 1i-l, Table S4). Finally, of the 3,611 SCOGs we had obtained for all the 54 samples, *Ranunculus sceleratus* (T-sample) had the most (3,453) and *Avena fatua* (T-sample) had the least (539) (Table S4). Also, the average number of SCOGs was higher in SD- samples (3,252) than in LN-samples (3,085) and significantly greater than in T-samples (2,939) (Figure 1h and l, Figure 2, Tables S2 and S3).

**Figure 2.**
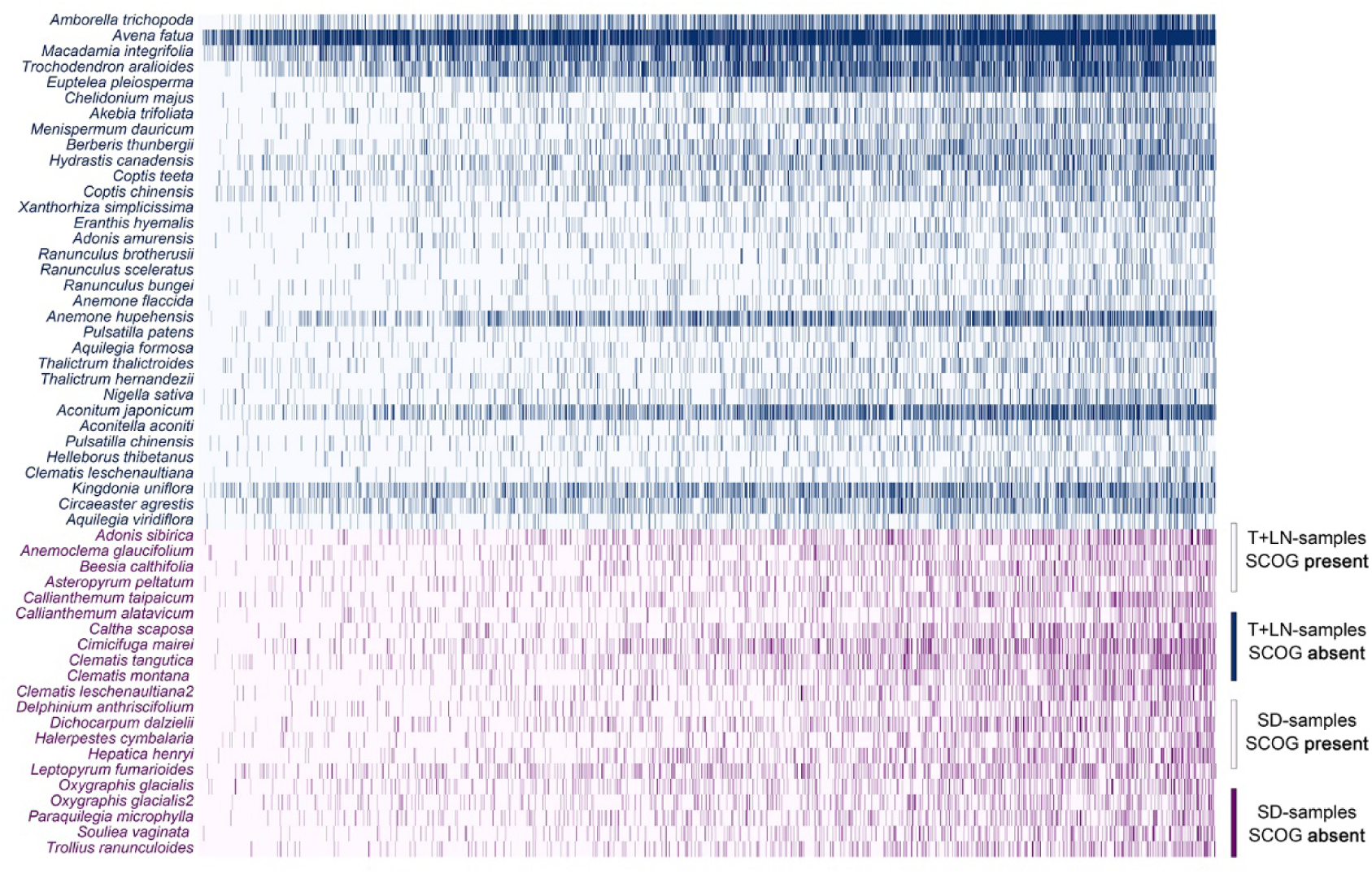
Heatmap of gene recovery efficiency. Each row represents one sample and each column represents one gene (3,611 genes total). Shading indicates a gene’s absence. The blue zone shows traditional treatment samples (i.e., tissues preserved in liquid nitrogen or in RNA stabilization solution) (see Table 1). The purple zone shows the 21 silica-dried samples from the current study

### 3.2 Phylogenetic inference

The ML analysis of the 3,611 SCOG-concatenated data set produced a highly resolved Ranunculaceae phylogeny in which all clades, except Caltheae + Nigelleae and Delphininieae + Cimicifugeae, were supported by 100 bootstrap values (Figure 3a). The 3,611 SCOG-coalescent-based analysis yielded a phylogeny very similar to the concatenated one, with high resolution and local posterior probability support (LPP, Figure 3b). However, both methods failed to confidently resolve *Nigella*’s position within the R-type clade. The other six more rigorously filtered data sets rendered phylogenetic frameworks that were completely congruent with the two 3,611 SCOG datasets (Figure S1). We then used the topology of the 3,611 SCOG-coalescent-based phylogeny (species tree) to discuss Ranunculaceae phylogeny and to conduct further analyses as the topology constraint.

**Figure 3.**
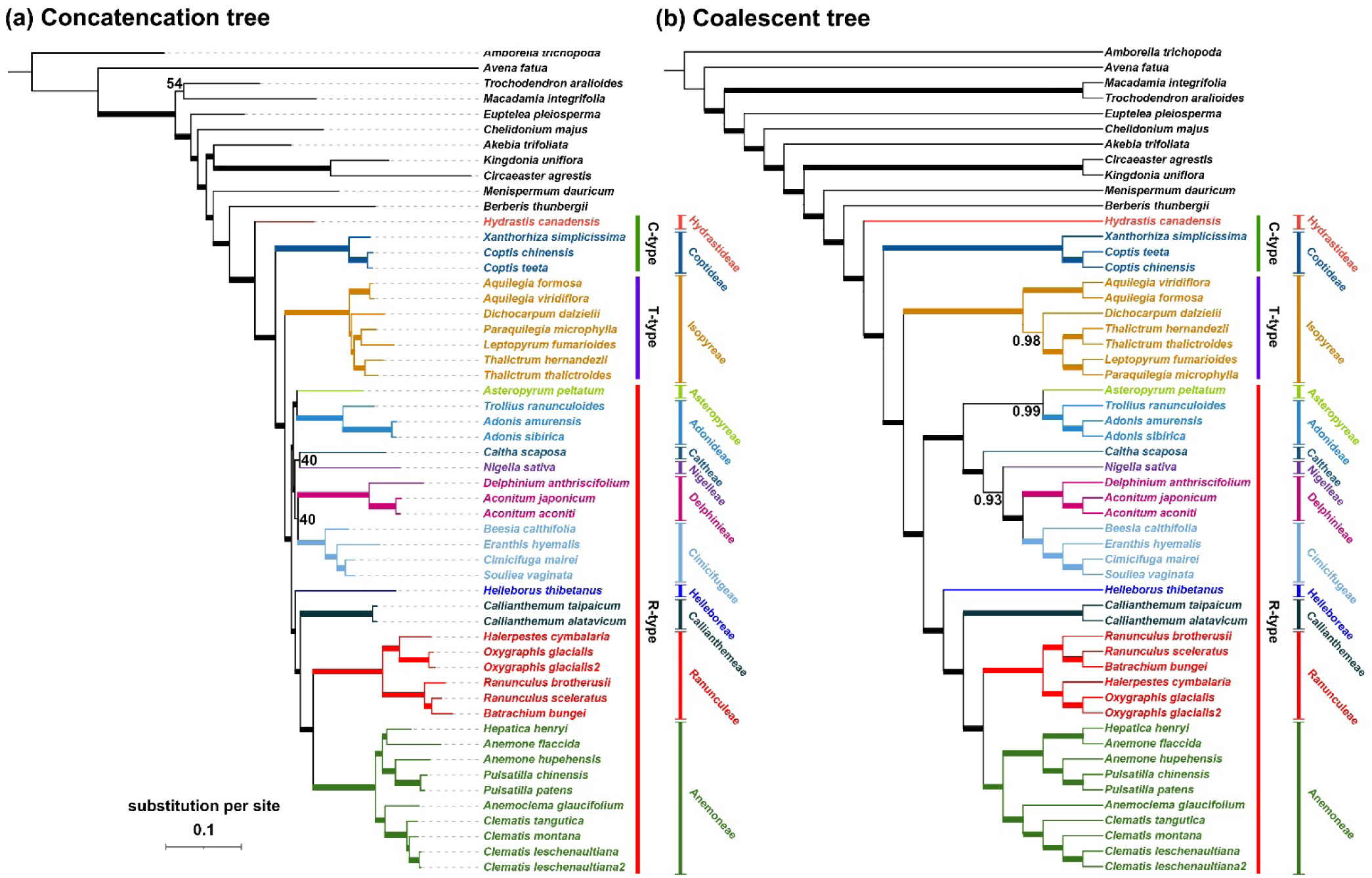
Phylogenetic relationships of Ranunculaceae. (a) The phylogeny inferred by the concatenated 3,611 single-copy co-orthologous nuclear gene (SCOG) data was produced using the maximum likelihood (ML) method. Bootstrap percentages are indicated on the branches and bold branches showed ML bootstrap values of 100. (b) The coalescence-based species tree topology was inferred by ASTRAL (Zhang, et al., 2018) from 3,611 nuclear genes. Numbers at branches are local posterior probabilities (ASTRAL-pp), but bold branches mark local posterior probabilities equal to 1.00. C-, T-, and R-types are chromosome groups and well-supported clades. The species in black are outgroups

The coalescent-based phylogeny (Figure 3b) resolved the 13 Ranunculaceae tribes we sampled (the first diverged Glaucideae was not available for this study), with the result consistent with previous studies (Wang et al., 2009, 2016; Zhai et al., 2019). Of those tribes, the Hydrastideae lineage diverged first within Ranunculaceae, followed by Coptideae (LPP = 1). The remaining eleven tribes (the core Ranunculaceae clade) formed two well supported major clades: T-type and R-type. The T-type clade included Isopyreae, in which all the genera have T-type chromosomes. The R-type clade included the remaining ten tribes whose genera have R-type chromosomes. There are two sub-clades within the R-type clade. One includes Helleboreae, Callianthemeae, Ranunculeae, and Anemoneae (LPP = 1) and the other contains the remaining six tribes. In the latter subclade, Asteropyreae and Adonideae are sister groups (LPP = 0.99), Caltheae is sister to Nigelleae, and the Caltheae – Nigelleae clade is then sister to the Delphinieae -Cimicifugeae clade.

### 3.3 Divergence time estimation for the Ranunculaceae

Dating results showed that the Ranunculaceae and Berberidaceae diverged in the Cretaceous, about 90.1 Mya with 95% highest posterior density interval (HPD) of 81.2–97.8 Mya (Figure S2). The crown age of the Ranunculaceae was estimated to be 80.2 Mya (95% HPD: 70.8–88.8 Mya) (Figure 4, Figure S2) and then the Coptideae clade diverged at about 71.7 Mya (95% HPD: 62.6–80.7 Mya). The T- and R-type clades diverged close to the Cretaceous-Paleogene (K-Pg) boundary at 66.7 Mya (95% HPD: 58.1–75.9 Mya) and the ten R-type tribes diversified in the early to middle Paleogene, from about 63 to 50 Mya.

**Figure 4.**
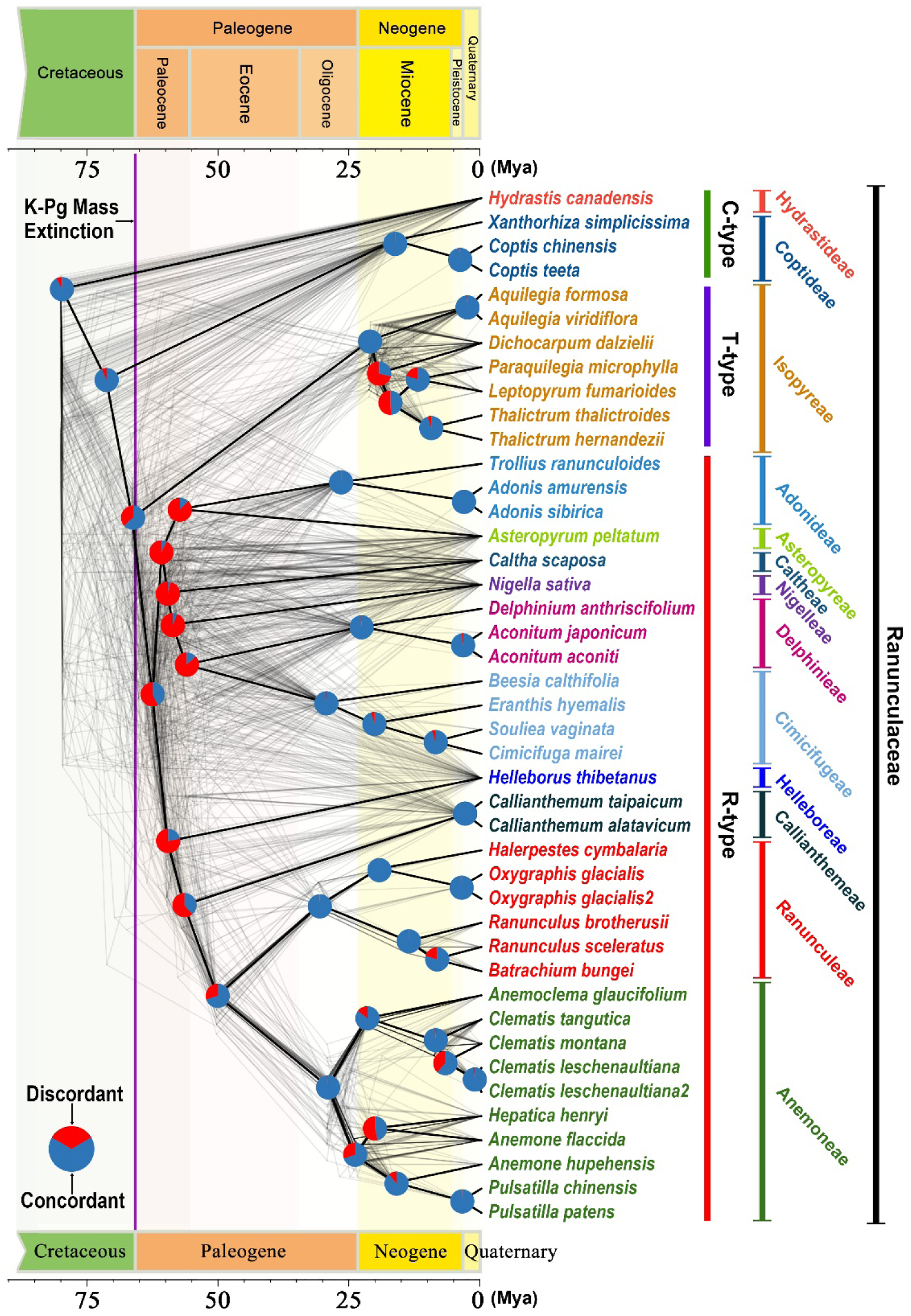
Coalescence-based species tree (Figure 3b) topology showing discordance among genes. The ASTRAL (Zhang et al., 2018) Ranunculaceae tree has all nodes resolved in most of the species trees (in heavy black lines). The gray-colored trees were sampled from 94 single-copy co-orthologous nuclear genes (from the 3,611 gene data set in which all the taxa were present in a single gene tree) and constructed with RAxML v.8.2.12. Pie charts on the nodes show proportions of concordant and discordant tree topologies compared to the species tree. C-, T-, and R-types are chromosome groups and well-supported clade

### 3.4 Conflict analyses

The conflicts between nuclear gene trees, as assessed by Phyparts v.0.0.1 showed a high proportion of concordant gene trees with the species tree, thus confirming the monophyly of the Ranunculaceae tribes that contain more than one genus (Figure 4). For example, Isopyreae (T-type clade) was supported by 97% (1,969/2,025) concordant, informative nuclear gene trees; Adonideae was supported by 1,609 of 1,705 (97%) informative gene trees; Ranunculeae was supported by 1,973 of 1,994 (99%) informative gene trees; and Anemoneae was supported by 2,037 of 2,068 (99%) informative gene trees. However, tribal relationships (nodes in Paleocene and early Eocene) were characterized by high levels of gene tree discordance (Figure 4).

We also compared the Ranunculaceae nuclear and plastid phylogenies (Figure 5) and found that the plastid phylogeny, having low support values at several nodes, was not better resolved than the nuclear phylogeny. Otherwise, the two phylogenies were largely consistent. The only supported cyto-nuclear discordance is the position of Adonineae (R-type). In the nuclear phylogeny, Adonieae is well-resolved in the R-type clade, but in the plastid phylogeny it is sister to Isopyreae in the T-type clade.

**Figure 5.**
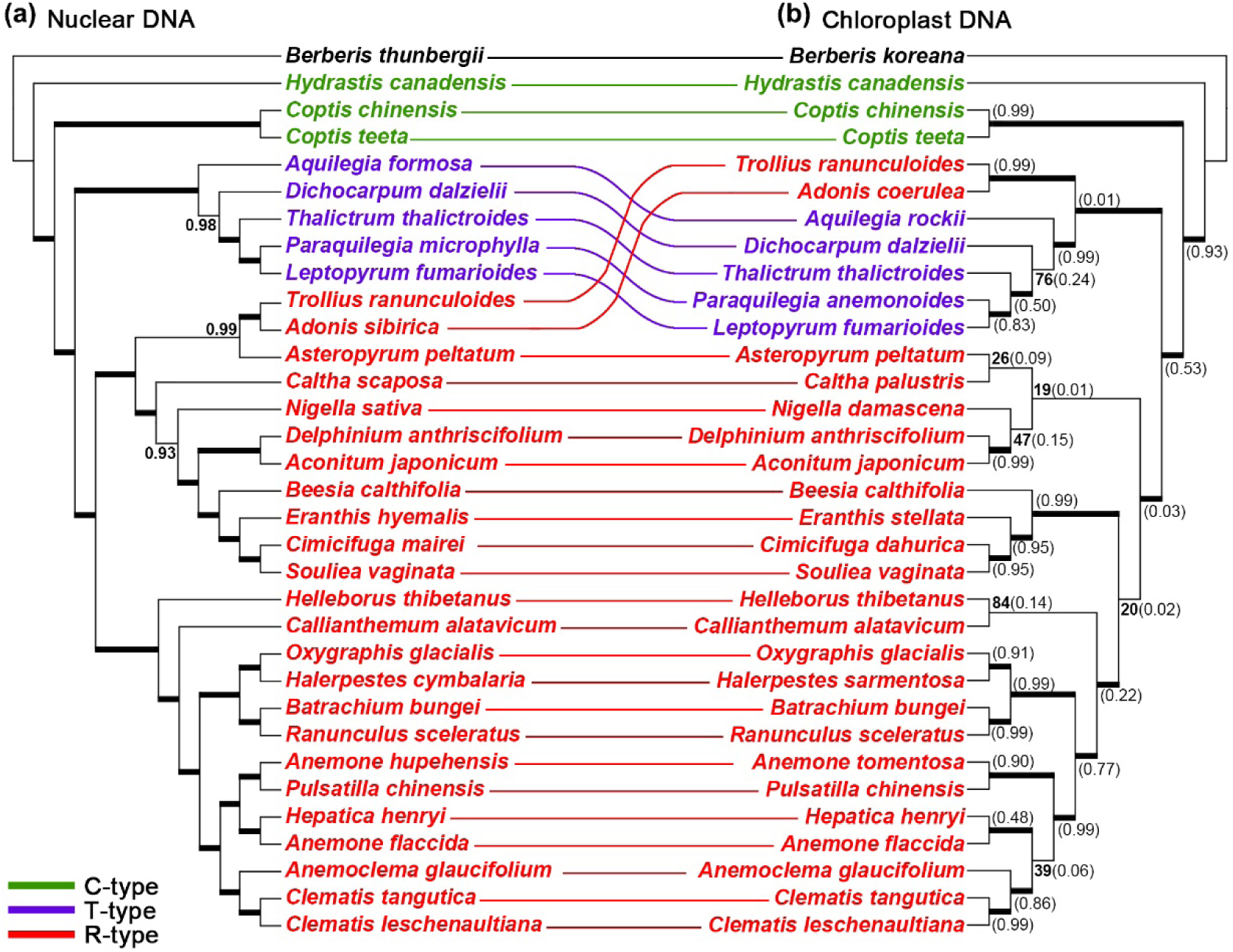
Comparison of the Ranunculaceae nuclear and plastid phylogenies. Left: Coalescence-based species tree topology inferred by ASTRAL (Zhang et al., 2018) from 3,611 nuclear genes (see Figure 3b). Numbers at branches are local posterior probabilities (ASTRAL-pp) but local posterior probabilities equal to 1.00 are marked with bold branches. Right: The Ranunculaceae phylogeny inferred from the concatenated coding regions of the complete plastome sequences using the maximum likelihood (ML) method. Bootstrap percentages are indicated on the branches and bold branch indicates ML bootstrap values of 100%. Numbers in brackets show the contribution of incomplete lineage sorting to the conflicts between the nuclear and plastid gene trees based on the multispecies coalescent model implemented in Phybase v.1.5 (Liu and Yu, 2010). Plastome sequence data are from previous studies (see Table S1)

### 3.5 Coalescent simulations and phylogenetic network estimating

To explain the cyto-nuclear discordance, we compared conflicts among the Phybase-simulated gene trees with the plastid genome phylogeny. The results showed that because of a very low proportion (0.01) of concordant simulated trees, the well-supported contradiction of the Adonineae between the nuclear and plastid trees could not be explained by ILS (Figure 5).

Nuclear gene discordance analysis (Figure 6a) showed much overlap between the empirical and simulated distance distributions, suggesting that ILS alone can explain most of the observed gene tree heterogeneity (Maureira-Butler et al., 2008). Meanwhile, during SNaQ analyses in PhyloNetworks, a plot of pseudo-loglikelihood scores (Figure 6c) suggested that the best network model was one with a maximum of one hybridization event (-ploglik = 246.7). We detected the most likely reticulation event to be between the *Caltha* and the ((*Callianthemum*, (*Ranunculus*, *Anemone*)), *Helleborus*) clades (γ = 0.368) with bootstrap value (BS) of 44.1 (Figure 6b). Also, when more than one hybridization event was allowed, we detected two additional reticulations in the Ranunculaceae species trees (Figure 6c), but the γ values of those two were minor (0.006–0.09).

**Figure 6.**
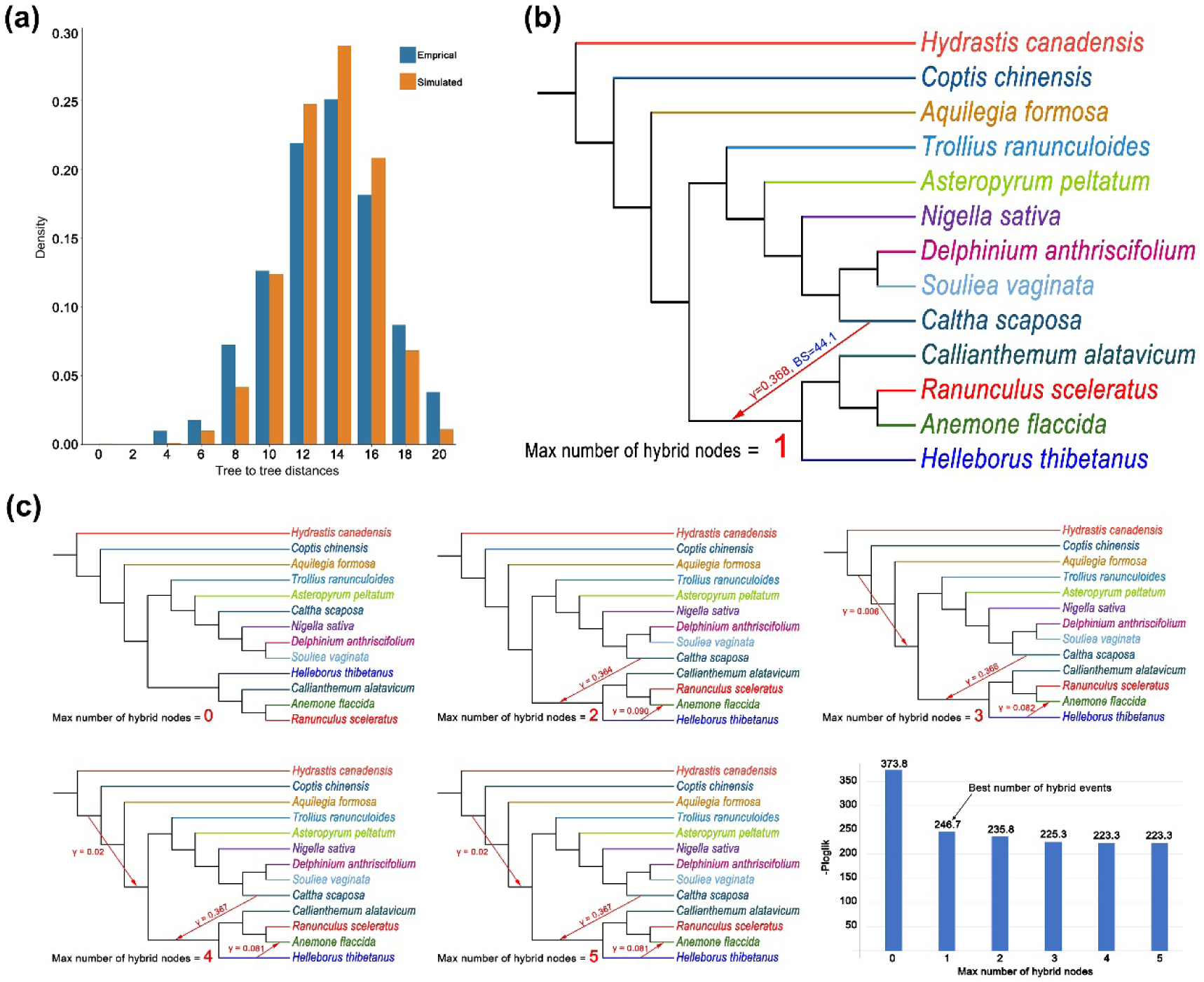
Testing the contributions of incomplete lineage sorting (ILS) and hybridization to the conflicts between the Ranunculaceae gene trees and the species tree inferred by ASTRAL (Zhang et al., 2018). (a) Coalescent simulations of tree-to-tree distance distributions between the empirical gene trees and the ASTRAL species tree (Empirical) and those from the coalescent simulation (Simulated). (b) Introgression model of unstable Ranunculaceae clades estimated using Phylonetworks. The results display maximum pseudolikelihood trees with a maximum zero to five allowed reticulations, and (c) the best model had a maximum allowed hybrid node of one according to the pseudo-loglikelihood scores (-ploglik) shown in the bar chart in (b). Red arrows mark possible reticulation events, γ is the gene contribution, and BS is the bootstrap support for the placement of minor hybrid branches

### 3.6 WGD detection

The results of Phyparts analysis showed a markedly increased number of gene duplications in several clades (Figure 7a), including the Ranunculales clade (R clade, 92 gene trees with duplication), the Ranunculaceae clade without Hydrastideae (RH clade, 86 gene trees with duplication), and the Delphinieae clade (DEL clade, 86 gene trees with duplication). We also found several clades with many gene trees with duplication (Figure 7a): the clade with all the Ranunculales families except Eupteleaceae (RE clade, 72 gene trees with duplication), the core Ranunculaceae clade (CR clade, 73 gene trees with duplication), and Anemoneae (ANE clade, 64 gene trees with duplication).

**Figure 7.**
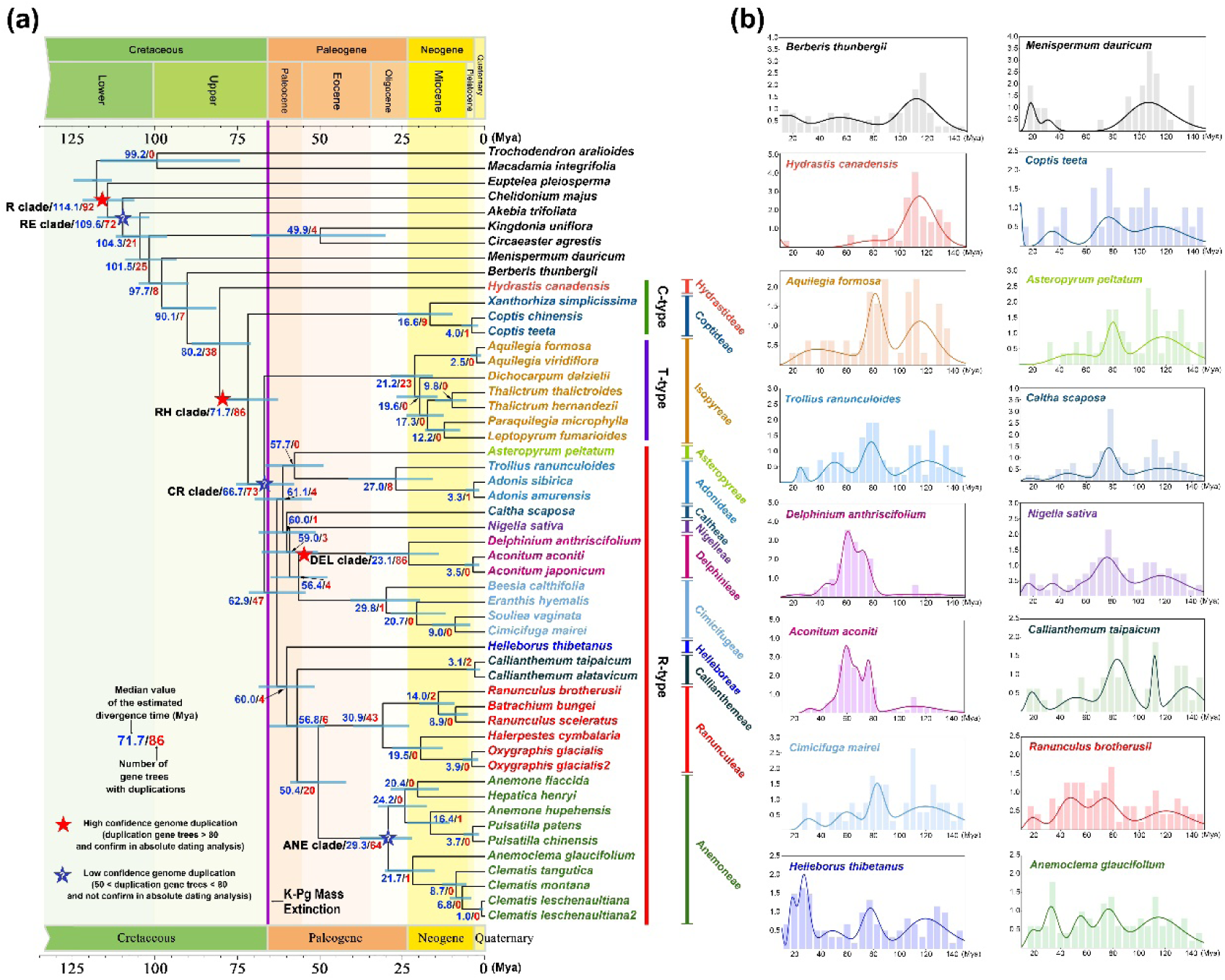
Genome duplication analysis under the absolute dating context. (a) Chronogram showing estimated divergence times according to MCMCTree and based on the Ranunculaceae coalescent species tree using 3,611 single-copy orthologous genes. The lineages have different colors and the taxa in a given lineage are grouped in one color. The outgroups are in black. Divergence times and numbers of gene trees with duplications (estimated by Phyparts) are marked under (or above) the branches. Starred clades are R: all Ranunculales, RE: R without Eupteleaceae, RH: R without Hydrastideae, CR: core Ranunculaceae, DEL: CR without Delphinieae, and ANE: CR without Anemoneae. (b) Absolute age distributions of genome duplications among major Ranunculales clades

Our absolute dating analysis (Figure 7b) found an obvious peak at around 80 Mya for each RH clade sample except *Hydrastis canadensis*, thus suggesting an ancient WGD event for this clade. We detected another pronounced peak at about 60 Mya for the DEL clade samples, suggesting an ancient WGD event for Delphinieae. We also detected a peak at about 110 Mya in some Ranunculaceae samples (e.g., *Hydrastis*, *Aquilegia*, *Callianthemum*, and *Helleborus*) and in the outgroups, and that may represent an ancient WGD event for Ranunculales. However, a few Ranunculaceae samples (e.g., *Aconitum* and *Caltha*), had no such peak. It is most likely because that later duplication may have relieved the selection pressure on gene copies and caused previously maintained copies to disappear (Ren et al., 2018). Those three WGD events corresponded with the three clades possessing the highest number of gene trees with duplication, as inferred by Phyparts analysis.

## 4 DISCUSSION

### 4.1 On using silica gel-dried leaf tissues in phylotranscriptomic analyses

For transcriptome partition, previous studies suggested that the contents and concentrations of total RNA, RIN, and rRNA ratio are the most important indices for assessing the success of RNA extraction (Jordon-Thaden et al., 2015; Yang et al., 2017). For example, an extraction was considered successful if the total RNA content was ≥ 30 μg, the RIN was higher than 5, and the rRNA ratio (26S/18S) was greater than 1 (Jordon-Thaden et al., 2015). Yang et al. (2017) considered an RIN higher than 6 and a concentration higher than 20 ng/μL acceptable. However, those indices may not be comparable among different studies because the acceptable RNA quality and quantity are greatly dependent on the next-generation sequencing platform used (Johnson et al., 2012). Nevertheless, our RNA extraction results showed that all the RIN values from the SD-samples were higher than six (average RIN = 7.5, Table S2). Although, most of those SD-samples did not yield ≥ 30 μg total RNA, the concentration of the extracted RNA was higher than 131.3 ng/μL (Table S2). Also, we found no significant differences in those RNA indices between the SD and LN treatments (Table S2).

Our results also showed that RNA does not decay as easily as previously thought, at least not in our Ranunculales samples. All 21 SD-samples yielded usable total RNA and among them, the SD-sample of *Clematis leschenaultiana* was stored in the -20 ℃ freezer for more than two years (Table 1, Table S2). Although SD-samples generated relatively lower quality transcriptome reads than the LN- and T-samples, and may not be useful for evo-devo studies, they are of sufficient quality for phylotranscriptomic analysis. Indeed, the assembled SCOGs from our SD-samples produced good or even better results than did those from the LN-samples and the T-samples used by the One Thousand Plant Transcriptomes Initiative (2019). If silica gel dried sampled can be used for RNA-seq in most angiosperm taxa, the utility of transcriptome data could become much more feasible for phylogenetic analyses andmay greatly improve the efficiency of data generation and reduce the cost. This SD plant tissue preservation strategy tested here provides novel insight into the feasibility of methods to easily obtain RNA datafor researchers in plant phylogenomics and may stimulate the exploration into broader groups of plants using SD preservation for RNA-seq phylogenetics.

### 4.2 Phylogenetic framework of the Ranunculaceae

Using Sanger sequencing data, molecular phylogenetic studies have not satisfactorily resolved the relationships of the 11 tribes that comprise the core Ranunculaceae (Johansson and Jansen, 1993; Hoot, 1995; Johansson, 1995; Ro et al., 1997; Hoot et al., 2008; Wang et al., 2009, 2016). Two chromosome types (R-type and T-type) are recognized in the core Rasnunculaceae (Tamura, 1995; Wang et al., 2009, 2016; Zhai et al., 2019).

Based on our 3,611 nuclear SCOGs (as well as the other rigorously selected data sets), we showed that the T- and R-type clades are robustly resolved as sister to each other within the core Ranunculaceae. According to our nuclear phylogeny, the T- and R- type chromosome groups seemed to have diverged from the C-type chromosome group. Our results are consistent with those of other published Ranunculaceae phylogenetic studies (Hoot, 1995; Hoot et al., 2008), and specifically support the Ranunculaceae phylogenetic classification proposed by Wang et al. (2009).

However, complete plastid genome data resolved different relationships within the core Ranunculaceae clade (Zhai et al., 2019; He et al., 2019). The sister relationship of Adonideae (R-type chromosome group) and Isopyreae (T-type chromosome group) were resolved and supported by the plastome sequences, while the other nine R-type tribes formed another clade and was sister to the Adonideae-Isopyreae clade (Zhai et al., 2019, Figure 5). This result indicated that T-type chromosomes may have derived from R-type chromosomes (Zhai et al., 2019). Strongly supported cyto-nuclear discordance may be the result of ILS or hybridization (Roos et al., 2011), but our coalescent simulating analysis suggests that ILS can not explain the cyto-nuclear discordance in Ranunculaceae. The different positions of Adonideae in the nuclear versus the plastid phylogenies may have been caused by ancient hybridization and/or subsequent introgression events, which are known to have happened in many angiosperm lineages (Liu et al., 2020; Hodel et al., 2021; Wang et al., 2021).

Within the R-type clade, the position of Cimicifugeae created another cyto-nuclear discordance. In our species tree (Figure 3b), Cimicifugeae was clustered with strong support as the sister group of Delphinieae. However, in the plastid phylogeny (Figure 5 and Zhai et al., 2019) Cimicifugeae was sister to a large R-type sub-clade including Helleboreae, Callianthemeae, Ranunculeae, and Anemoneae, but it had low support values. This discordance was likely caused by insufficient information in the plastid genome data.

Using complete plastid genome data, Zhai et al. (2019) estimated two waves of radiation in the Ranunculaceae after the Cretaceous-Paleogene (K-P) boundary. Our molecular dating analysis yielded similar divergence time estimates and consistently revealed two waves of radiation after the K-P boundary (Figure 4). Almost all the ten tribes of the R-type clade evolved in the first wave of radiation during the Paleocene and early Eocene eras. Furthermore, we detected widespread nuclear gene discordance during that diversification, thus showing complicated evolutionary processes at work during that period. Our coalescent simulating results showed that the conflict among nuclear trees was mainly due to ancestral genetic polymorphism and ILS, and supported one hybridization event between Caltheae and the (Callianthemeae-[Ranunculeae-Anemoneae]-Helleboreae) clades (Figure 6). Our results have hence provided important insights into why previous phylogenetic studies, based on either limited numbers of genes or uniparentally transmitted markers, could not satisfactorily resolve the tribal relationships of Ranunculaceae.

### 4.3 Whole genome duplications

Our WGD analyses support three WGD events within the Ranunculales (Figure 7). The first event occurred in the Ranunculales ancestor at around 110 Mya, as previous studies had also suggested (Pabón-Mora et al., 2013; One Thousand Plant Transcriptomes Initiatives, 2019; Pei et al., 2021). After that, we detected another WGD event around 80 Mya after the divergence of Hydrastideae, but before the divergence of Coptideae and the core Ranunculaceae clade. Previous studies also reported an ancient WGD event either after the divergence of Berberidaceae and Ranunculaceae (Xie et al., 2020) or shared by the Ranunculaceae (Liu et al., 2021). However, those studies did not include samples of the Ranunculaceae tribes that diverged the earliest, the Glaucideae and Hydrastideae. So, those reported WGDs are likely the second WGD event that we detected before the divergence of the Coptideae and the core Ranunculaceae clade. A third WGD event was detected in the stem lineage of Delphinieae by Park et al. (2020) who, using plastid genome and transcriptome data, found that plastid genes *rpl*32 and *rps*16 were transferred to the nucleus in Delphinieae and duplicated in *Aconitum* samples. They assumed that this duplication was associated with an ancient WGD event in *Aconitum*. However, the WGD hypothesis based on the two duplicated genes needs to be tested with more robust data. Our results suggest that both *Aconitum* and *Delphinium* may have shared an ancient WGD event. Both the CR and R-type clades had large numbers of gene trees with gene duplication (73 for the CR clade and 47 for the R-type clade, Figure 7a), but the dating analysis did not support the WGD events in these two clades. The high gene duplication detected in those clades may have been caused by small scale or single gene duplications or other mechanisms.

### 4.4 Conclusions

In this study, we tested whether silica gel-dried plant tissues could be used for RNA extraction and phylotranscriptomic studies. Our results showed that the RNA extracted from silica gel-dried leaf tissue samples of members of the Ranunculaceae and the close relatives performed well in subsequent phylogenetic analyses. Based on the transcriptome data we generated, we successfully resolved long-standing phylogenetic controversies in the Ranunculaceae, such as core Ranunculaceae tribal relationships, and cyto-nuclear discordance of the T- and R-type clades. We also obtained a resolved Ranunculaceae phylogeny that had better statistical support than in previous studies. The 11 tribes of the core Ranunculaceae grouped into two clades corresponding to the chromosome types (the T- and R-type clades), with high support. Two waves of radiation, previously inferred based on plastome data (Zhai et al., 2019), were also supported by our nuclear SCOG data. The first wave, which represented the deep diversification of all 11 tribes of the core Ranunculaceae, underwent extremely complicated evolutionary processes, such as gene duplication, hybridization, introgression, and incomplete lineage sorting. However, WGD was important in the evolution of Ranunculaceae, but not a direct factor in chromosome type evolution in the family. Our results indicated that using either a limited number of genes or plastid genome data does not satisfactorily resolve Ranunculaceae tribal relationships. Our results provide novel insights into RNA-seq methods that may greatly increase the utility of RNA sequencing in phylogenomics, as well as into the phylogenetics and evolution of the widely contested Ranunculaceae family.

## ACKNOWLEDGEMENTS

This study was supported by the National Natural Science Foundation of China [grant number 31670207] and the Beijing Natural Science Foundation [grant number 5182016].

## DATA AVAILABILITY STATEMENT

Raw sequence reads are deposited in the NCBI SRA BioProject ID PRJNA739247; Tree, alignment and code are deposited in the Zenodo (https://doi.org/10.5281/zenodo.5144194).

## AUTHOR CONTRIBUTIONS

Jian He: Formal analysis, Methodology, Software, Validation, Visualization, Writing original draft. Rudan Lyu: Investigation, Resources, Data curation. Yike Luo: Investigation, Resources. Jiamin Xiao: Investigation, Data curation. Lei Xie: Writing - review & editing, Funding acquisition. Jun Wen: Writing - review & editing, Supervision. Wenhe Li: Investigation, Resources. Linying Pei: Investigation, Resources. Jin Cheng: Project administration, Supervision.

## SUPPLEMENTAL INFORMATION

**SUPPLEMENTARY MATERIALS** Protocol of RNA extraction used in this study

FIGURE S1 Ranunculaceae phylogenies inferred from nuclear datasets with different selective constrains (see Materials and Methods, 2.3). Phylogenies inferred from concatenated datasets were produced using the maximum likelihood method. Bootstrap percentages are indicated on the branches and bold branch shows ML bootstrap values of 100. Coalescence-based species tree topologies were inferred by ASTRAL (Zhang et al., 2018). Numbers at branches are local posterior probabilities (ASTRAL-pp), but bold branches mark local posterior probabilities equal to 1.00

FIGURE S2 Chronogram of Ranunculaceae inferred from the ASTRAL (Zhang et al., 2018) topology of the 3,611 SCOG coalescent data set using MCMCTree implemented in the PAML v.4.9a package (Yang, 2007). Red stars mark fossil constraints which are explained in the text (Materials and Methods 2.5). Numbers on the branches indicate the median value (95% high posterior density interval) of the estimated divergence time (Mya)

TABLE S1 Species sampling information and references for plastome sequences used in Figure 5

TABLE S2 RNA quality and sequencing information for the newly sequenced samples in this study. One-way ANOVAs for each column are included in worksheets in this Excel workbook

TABLE S3 Information of de novo assembly of newly sequenced samples in this study. One-way ANOVAs for each column are included in worksheets in this Excel workbook

TABLE S4 Information about the de novo assembly of all samples (including newly sequenced and downloaded) in this study. One-way ANOVAs for each column are included in worksheets in this Excel workbook

