## SUPPLEMENTARY MATERIALS for "A phylotranscriptome study using silica gel-dried leaf tissues produces an updated robust phylogeny of Ranunculaceae" (from another version of this preprint: version 1, 2021-07-29)

1. Place silica-dried leaves in liquid nitrogen and grind thoroughly with a mortar and pestle, Use 1 ml TRNzol Universal Reagent per 100 mg leaves.
2. Incubate homogenized samples at 15-30°C for 5 min, to permit complete dissociation of the nucleoprotein complex.
3. Add 0.2 ml chloroform per 1 ml TRNzol Universal Reagent used for homogenization. Cap the tube securely and vortex for 15 sec. Incubate for 3 min at room temperature.
4. Centrifuge the sample at 12,000 rpm (~13,400 × g) for 15 min at 4°C. The mixture separates into a lower pink phenol-chloroform phase, an interphase, and a colorless upper aqueous phase. RNA remains exclusively in the aqueous phase. Pipette the aqueous phase (about 600 ul) out into a new tube.
5. Add equal volume of 100% isopropanol to the aqueous phase. Mix thoroughly and incubate at room temperature for 10 min.
6. Centrifuge at 12,000 rpm (~13,400 × g) for 10 min at 4°C. Remove the supernatant from the tube, leaving only the RNA pellet.
7. Wash the pellet, with at least 1 ml of 75% ethanol (in RNase-Free ddH_2_O) per 1 ml of TRNzol Universal Reagent used in the initial homogenization.
8. Centrifuge the tube at 10,000 rpm (~9,391 × g) for 5 min at 4°C. Discard the wash. Centrifuge briefly to collect residual liquid and pipette them out without drawing the pellet.
9. Air dry the RNA pellet for 2-3 min. (Do not allow the RNA to dry completely, because the pellet can lose solubility.) According to the requirement, add 30-100 μl RNase-Free ddH_2_O, resuspend the RNA pellet completely by passing the solution up and down repeated through a pipette tip.
