## Supplementary material for "A phylotranscriptome study using silica gel-dried leaf tissues produces an updated robust phylogeny of Ranunculaceae": FIGURE S1 (from another version of this preprint: version 1, 2021-07-29)

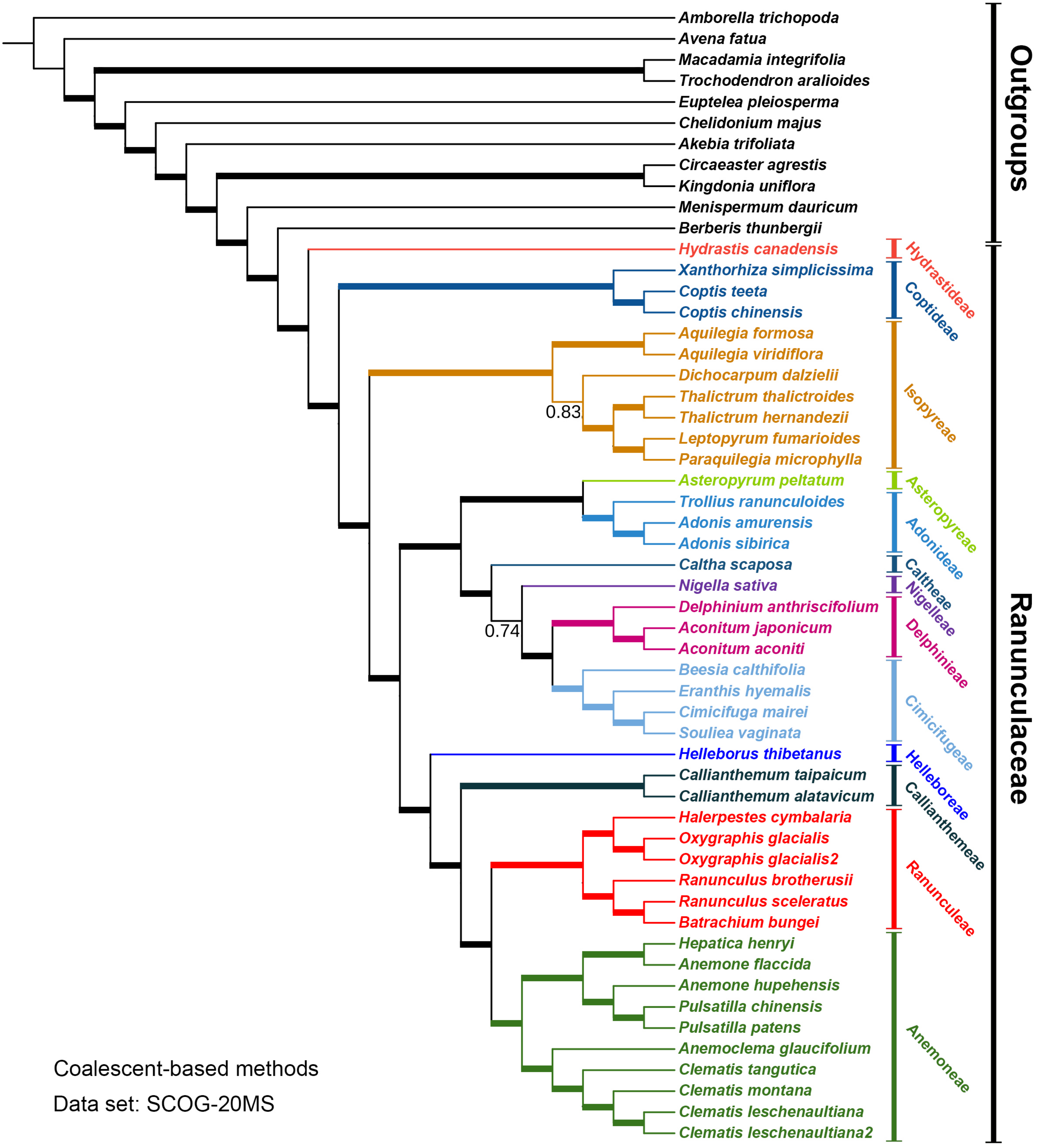

Amborella trichopoda
Avena fatua
Macadamia integrifolia
Trochodendron aralioides
Euptelea pleiosperma
Chelidonium majus
Akebia trifoliata
Circaeaster agrestis
Kingdonia uniflora
Menispermum dauricum
Berberis thunbergii
Hydrastis canadensis
Xanthorhiza simplicissima
Coptis teeta
Coptis chinensis
Aquilegia formosa
Aquilegia viridiflora
Dichocarpum dalzielii
Thalictrum thalictroides
Thalictrum hernandezii
Leptopyrum fumarioides
Paraquilegia microphylla
0.83
Asteropyrum peltatum
Trollius ranunculoides
Adonis amurensis
Adonis sibirica
Caltha scaposa
Nigella sativa
Delphinium anthriscifolium
Aconitum japonicum
Aconitum aconiti
0.74
Beesia calthifolia
Eranthis hyemalis
Cimicifuga mairei
Souliea vaginata
Helleborus thibetanus
Callianthemum taipaicum
Callianthemum alatavicum
Halerpestes cymbalaria
Oxygraphis glacialis
Oxygraphis glacialis2
Ranunculus brotherusii
Ranunculus sceleratus
Batrachium bungei
Hepatica henryi
Anemone flaccida
Anemone hupehensis
Pulsatilla chinensis
Pulsatilla patens
Anemoclema glaucifolium
Clematis tangutica
Clematis montana
Clematis leschenaultiana
Clematis leschenaultiana2
Outgroups
Ranunculaceae
Hydrastideae
Coptideae
Isopyreae
Asteropyreae
Adonideae
Caltheae
Nigelleae
Delphinieae
Cimicifugeae
Helleboreae
Callianthemeae
Ranunculeae
Anemoneae
Coalescent-based methods
Data set: SCOG-20MS

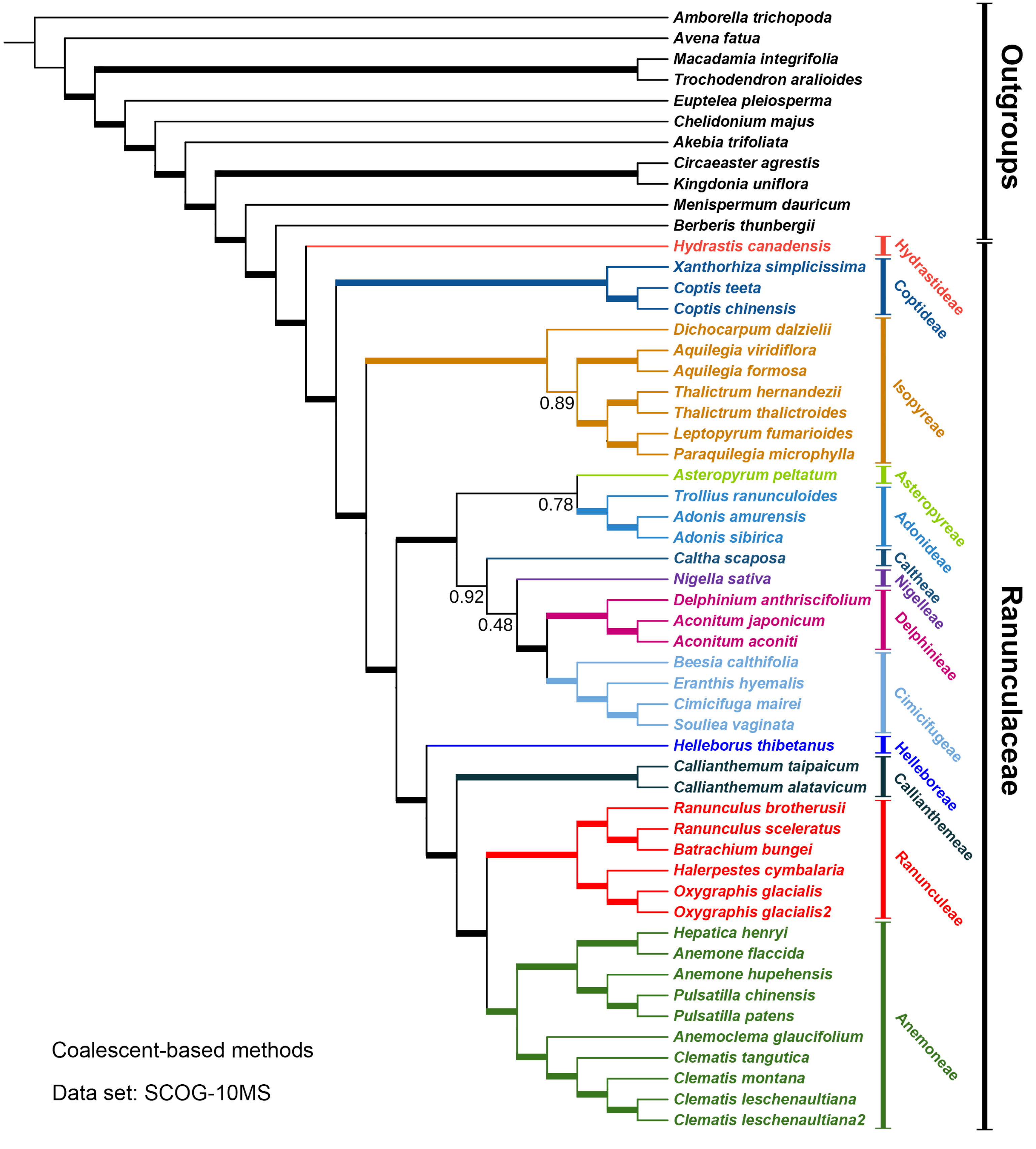

Amborella trichopoda
Avena fatua
Macadamia integrifolia
Trochodendron aralioides
Euptelea pleiosperma
Chelidonium majus
Akebia trifoliata
Circaeaster agrestis
Kingdonia uniflora
Menispermum dauricum
Berberis thunbergii
Hydrastis canadensis
Xanthorhiza simplicissima
Coptis teeta
Coptis chinensis
Dichocarpum dalzielii
Aquilegia viridiflora
Aquilegia formosa
Thalictrum hernandezii
Thalictrum thalictroides
Leptopyrum fumarioides
Paraquilegia microphylla
Asteropyrum peltatum
Trollius ranunculoides
Adonis amurensis
Adonis sibirica
Caltha scaposa
Nigella sativa
Delphinium anthriscifolium
Aconitum japonicum
Aconitum aconiti
Beesia calthifolia
Eranthis hyemalis
Cimicifuga mairei
Souliea vaginata
Helleborus thibetanus
Callianthemum taipaicum
Callianthemum alatavicum
Ranunculus brotherusii
Ranunculus sceleratus
Batrachium bungei
Halerpestes cymbalaria
Oxygraphis glacialis
Oxygraphis glacialis2
Hepatica henryi
Anemone flaccida
Anemone hupehensis
Pulsatilla chinensis
Pulsatilla patens
Anemoclema glaucifolium
Clematis tangutica
Clematis montana
Clematis leschenaultiana
Clematis leschenaultiana2
0.89
0.78
0.92
0.48
Hydrastideae
Coptideae
Isopyreae
Asteropyreae
Adonideae
Caltheae
Nigelleae
Delphinieae
Cimicifugeae
Helleboreae
Callianthemeae
Ranunculeae
Anemoneae
Outgroups
Ranunculaceae
Coalescent-based methods
Data set: SCOG-10MS

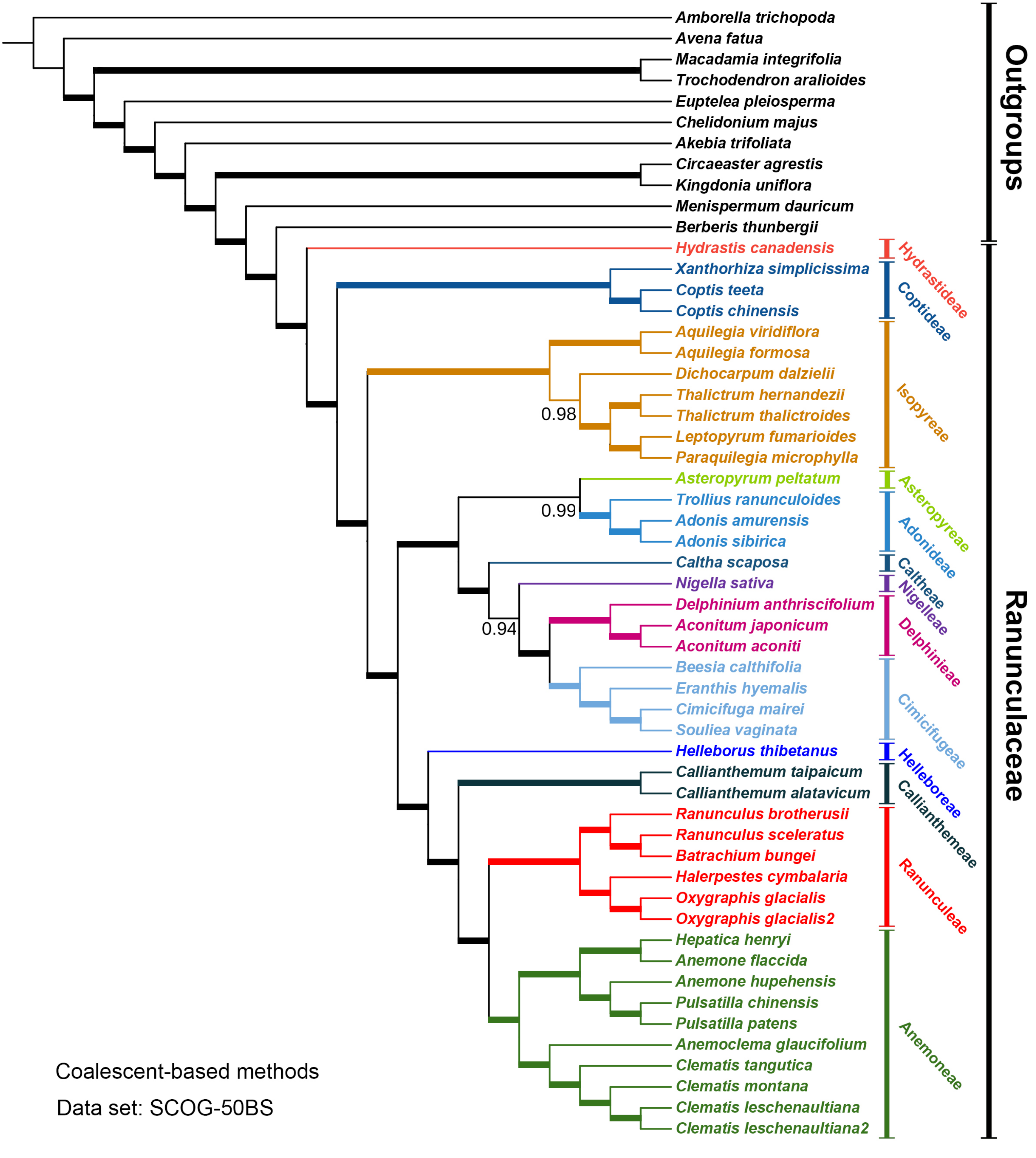

Amborella trichopoda
Avena fatua
Macadamia integrifolia
Trochodendron aralioides
Euptelea pleiosperma
Chelidonium majus
Akebia trifoliata
Circaeaster agrestis
Kingdonia uniflora
Menispermum dauricum
Berberis thunbergii
Hydrastis canadensis
Xanthorhiza simplicissima
Coptis teeta
Coptis chinensis
Aquilegia viridiflora
Aquilegia formosa
Dichocarpum dalzielii
Thalictrum hernandezii
Thalictrum thalictroides
Leptopyrum fumarioides
Paraquilegia microphylla
Asteropyrum peltatum
Trollius ranunculoides
Adonis amurensis
Adonis sibirica
Caltha scaposa
Nigella sativa
Delphinium anthriscifolium
Aconitum japonicum
Aconitum aconiti
Beesia calthifolia
Eranthis hyemalis
Cimicifuga mairei
Souliea vaginata
Helleborus thibetanus
Callianthemum taipaicum
Callianthemum alatavicum
Ranunculus brotherusii
Ranunculus sceleratus
Batrachium bungei
Halerpestes cymbalaria
Oxygraphis glacialis
Oxygraphis glacialis2
Hepatica henryi
Anemone flaccida
Anemone hupehensis
Pulsatilla chinensis
Pulsatilla patens
Anemoclema glaucifolium
Clematis tangutica
Clematis montana
Clematis leschenaultiana
Clematis leschenaultiana2
0.98
0.99
0.94
Outgroups
Ranunculaceae
Hydrastideae
Coptideae
Isopyreae
Asteropyreae
Adonideae
Caltheae
Nigelleae
Delphinieae
Cimicifugeae
Helleboreae
Callianthemeae
Ranunculeae
Anemoneae
Coalescent-based methods
Data set: SCOG-50BS

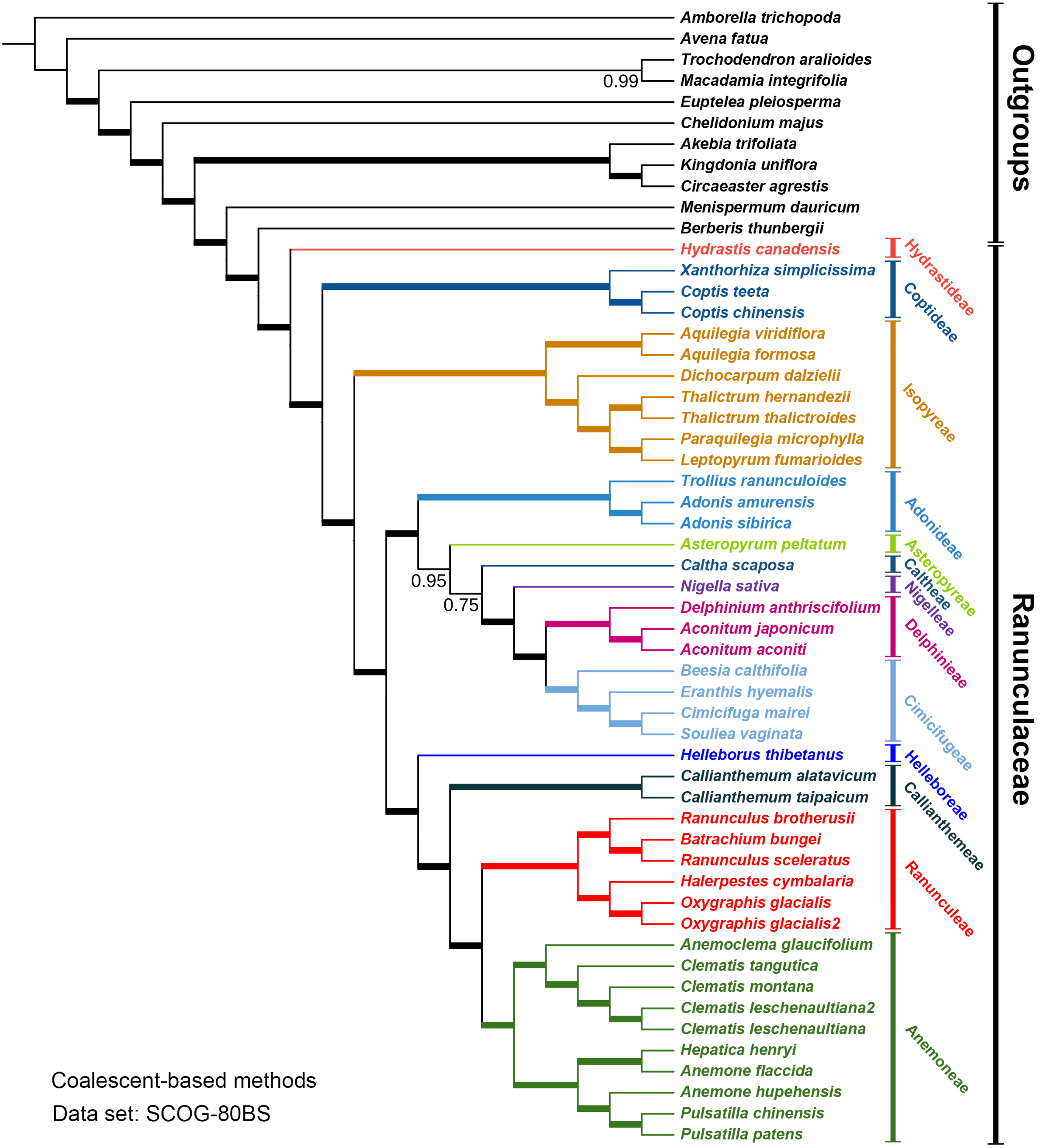

Amborella trichopoda
Avena fatua
Trochodendron aralioides
Macadamia integrifolia
0.99
Euptelea pleiosperma
Chelidonium majus
Akebia trifoliata
Kingdonia uniflora
Circaeaster agrestis
Menispermum dauricum
Berberis thunbergii
Hydrastis canadensis
Xanthorhiza simplicissima
Coptis teeta
Coptis chinensis
Aquilegia viridiflora
Aquilegia formosa
Dichocarpum dalzielii
Thalictrum hernandezii
Thalictrum thalictroides
Paraquilegia microphylla
Leptopyrum fumarioides
Trollius ranunculoides
Adonis amurensis
Adonis sibirica
Asteropyrum peltatum
Caltha scaposa
Nigella sativa
0.95
0.75
Delphinium anthriscifolium
Aconitum japonicum
Aconitum aconiti
Beesia calthifolia
Eranthis hyemalis
Cimicifuga mairei
Souliea vaginata
Helleborus thibetanus
Callianthemum alatavicum
Callianthemum taipaicum
Ranunculus brotherusii
Batrachium bungei
Ranunculus sceleratus
Halerpestes cymbalaria
Oxygraphis glacialis
Oxygraphis glacialis2
Anemoclema glaucifolium
Clematis tangutica
Clematis montana
Clematis leschenaultiana2
Clematis leschenaultiana
Hepatica henryi
Anemone flaccida
Anemone hupehensis
Pulsatilla chinensis
Pulsatilla patens
Hydrastideae
Coptideae
Isopyreae
Adonideae
Asteropyreae
Caltheae
Nigelleae
Delphinieae
Cimicifugeae
Helleboreae
Callianthemeae
Ranunculeae
Anemoneae
Outgroups
Ranunculaceae
Coalescent-based methods
Data set: SCOG-80BS

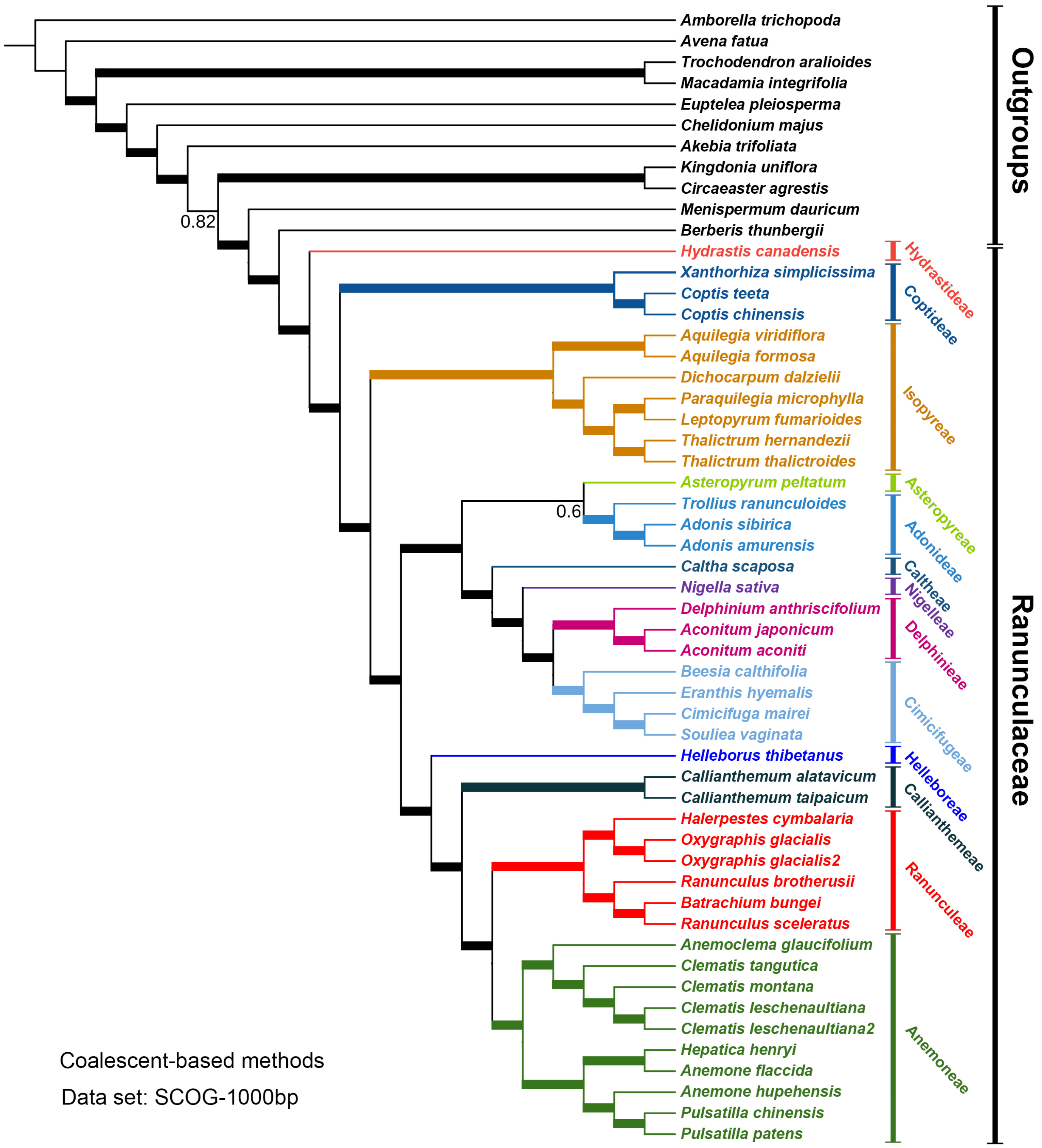
Amborella trichopoda
Avena fatua
Trochodendron aralioides
Macadamia integrifolia
Euptelea pleiosperma
Chelidonium majus
Akebia trifoliata
Kingdonia uniflora
Circaeaster agrestis
Menispermum dauricum
Berberis thunbergii
0.82
Hydrastis canadensis
Xanthorhiza simplicissima
Coptis teeta
Coptis chinensis
Aquilegia viridiflora
Aquilegia formosa
Dichocarpum dalzielii
Paraquilegia microphylla
Leptopyrum fumarioides
Thalictrum hernandezii
Thalictrum thalictroides
Asteropyrum peltatum
0.6
Trollius ranunculoides
Adonis sibirica
Adonis amurensis
Caltha scaposa
Nigella sativa
Delphinium anthriscifolium
Aconitum japonicum
Aconitum aconiti
Beesia calthifolia
Eranthis hyemalis
Cimicifuga mairei
Souliea vaginata
Helleborus thibetanus
Callianthemum alatavicum
Callianthemum taipaicum
Halerpestes cymbalaria
Oxygraphis glacialis
Oxygraphis glacialis2
Ranunculus brotherusii
Batrachium bungei
Ranunculus sceleratus
Anemoclema glaucifolium
Clematis tangutica
Clematis montana
Clematis leschenaultiana
Clematis leschenaultiana2
Hepatica henryi
Anemone flaccida
Anemone hupehensis
Pulsatilla chinensis
Pulsatilla patens
Outgroups
Ranunculaceae
Hydrastideae
Coptideae
Isopyreae
Asteropyreae
Adonideae
Caltheae
Nigelleae
Delphinieae
Cimicifugeae
Helleboreae
Callianthemeae
Ranunculeae
Anemoneae
Coalescent-based methods
Data set: SCOG-1000bp

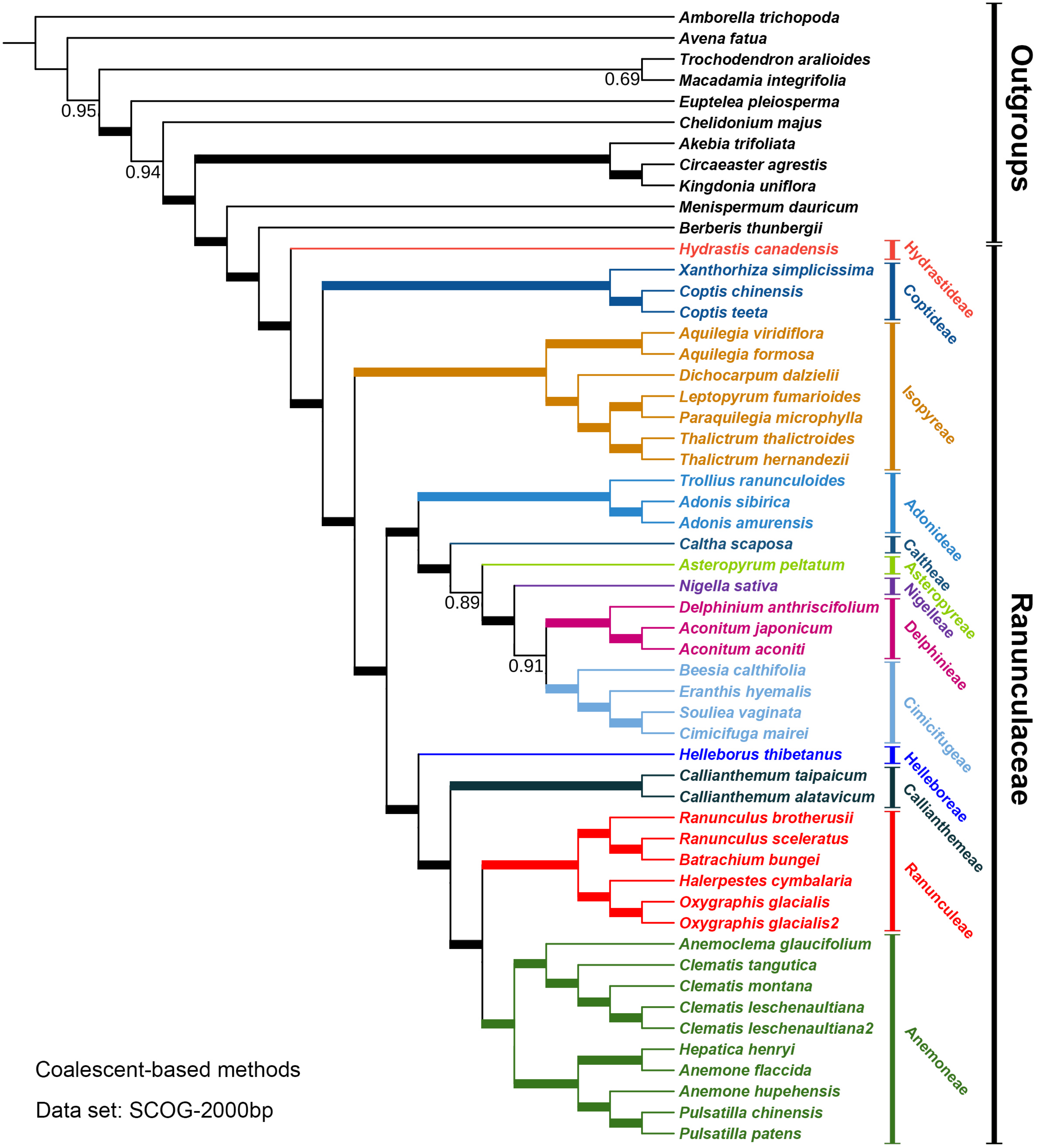
Amborella trichopoda
Avena fatua
Trochodendron aralioides
Macadamia integrifolia
Euptelea pleiosperma
Chelidonium majus
Akebia trifoliata
Circaeaster agrestis
Kingdonia uniflora
Menispermum dauricum
Berberis thunbergii
Hydrastis canadensis
Xanthorhiza simplicissima
Coptis chinensis
Coptis teeta
Aquilegia viridiflora
Aquilegia formosa
Dichocarpum dalzielii
Leptopyrum fumarioides
Paraquilegia microphylla
Thalictrum thalictroides
Thalictrum hernandezii
Trollius ranunculoides
Adonis sibirica
Adonis amurensis
Caltha scaposa
Asteropyrum peltatum
Nigella sativa
Delphinium anthriscifolium
Aconitum japonicum
Aconitum aconiti
Beesia calthifolia
Eranthis hyemalis
Souliea vaginata
Cimicifuga mairei
Helleborus thibetanus
Callianthemum taipaicum
Callianthemum alatavicum
Ranunculus brotherusii
Ranunculus sceleratus
Batrachium bungei
Halerpestes cymbalaria
Oxygraphis glacialis
Oxygraphis glacialis2
Anemoclema glaucifolium
Clematis tangutica
Clematis montana
Clematis leschenaultiana
Clematis leschenaultiana2
Hepatica henryi
Anemone flaccida
Anemone hupehensis
Pulsatilla chinensis
Pulsatilla patens
0.69
0.95
0.94
0.89
0.91
Outgroups
Ranunculaceae
Hydrastideae
Coptideae
Isopyreae
Adonideae
Caltheae
Asteropyreae
Nigelleae
Delphinieae
Cimicifugeae
Helleboreae
Callianthemeae
Ranunculeae
Anemoneae
Coalescent-based methods
Data set: SCOG-2000bp

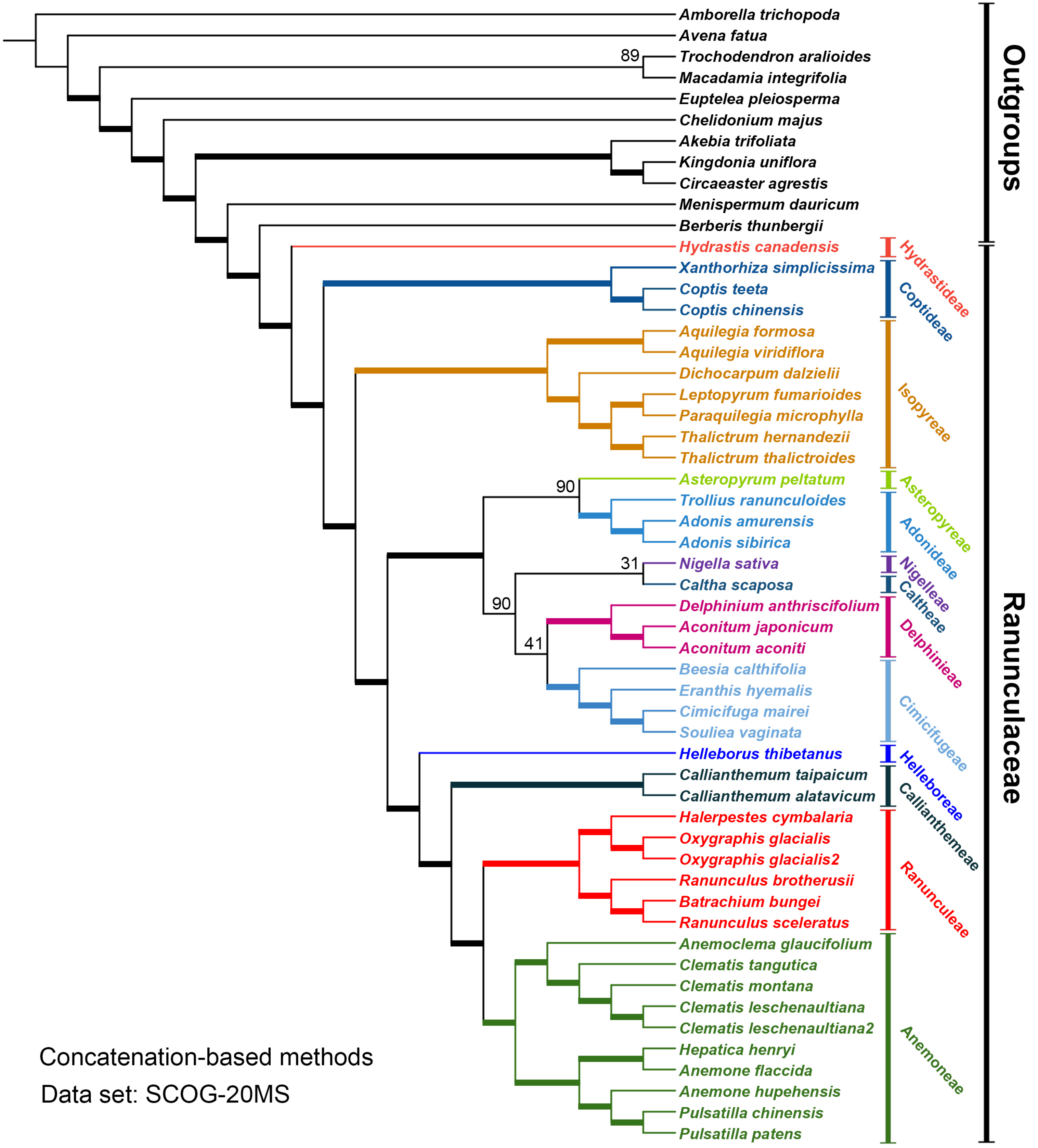
Amborella trichopoda
Avena fatua
89
Trochodendron aralioides
Macadamia integrifolia
Euptelea pleiosperma
Chelidonium majus
Akebia trifoliata
Kingdonia uniflora
Circaeaster agrestis
Menispermum dauricum
Berberis thunbergii
Hydrastis canadensis
Xanthorhiza simplicissima
Coptis teeta
Coptis chinensis
Aquilegia formosa
Aquilegia viridiflora
Dichocarpum dalzielii
Leptopyrum fumarioides
Paraquilegia microphylla
Thalictrum hernandezii
Thalictrum thalictroides
90
Asteropyrum peltatum
Trollius ranunculoides
Adonis amurensis
Adonis sibirica
31
Nigella sativa
Caltha scaposa
90
Delphinium anthriscifolium
Aconitum japonicum
41
Aconitum aconiti
Beesia calthifolia
Eranthis hyemalis
Cimicifuga mairei
Souliea vaginata
Helleborus thibetanus
Callianthemum taipaicum
Callianthemum alatavicum
Halerpestes cymbalaria
Oxygraphis glacialis
Oxygraphis glacialis2
Ranunculus brotherusii
Batrachium bungei
Ranunculus sceleratus
Anemoclema glaucifolium
Clematis tangutica
Clematis montana
Clematis leschenaultiana
Clematis leschenaultiana2
Hepatica henryi
Anemone flaccida
Anemone hupehensis
Pulsatilla chinensis
Pulsatilla patens
Outgroups
Hydrastideae
Coptideae
Isopyreae
Asteropyreae
Adonideae
Nigelleae
Caltheae
Delphinieae
Cimicifugeae
Helleboreae
Callianthemeae
Ranunculeae
Anemoneae
Ranunculaceae
Concatenation-based methods
Data set: SCOG-20MS

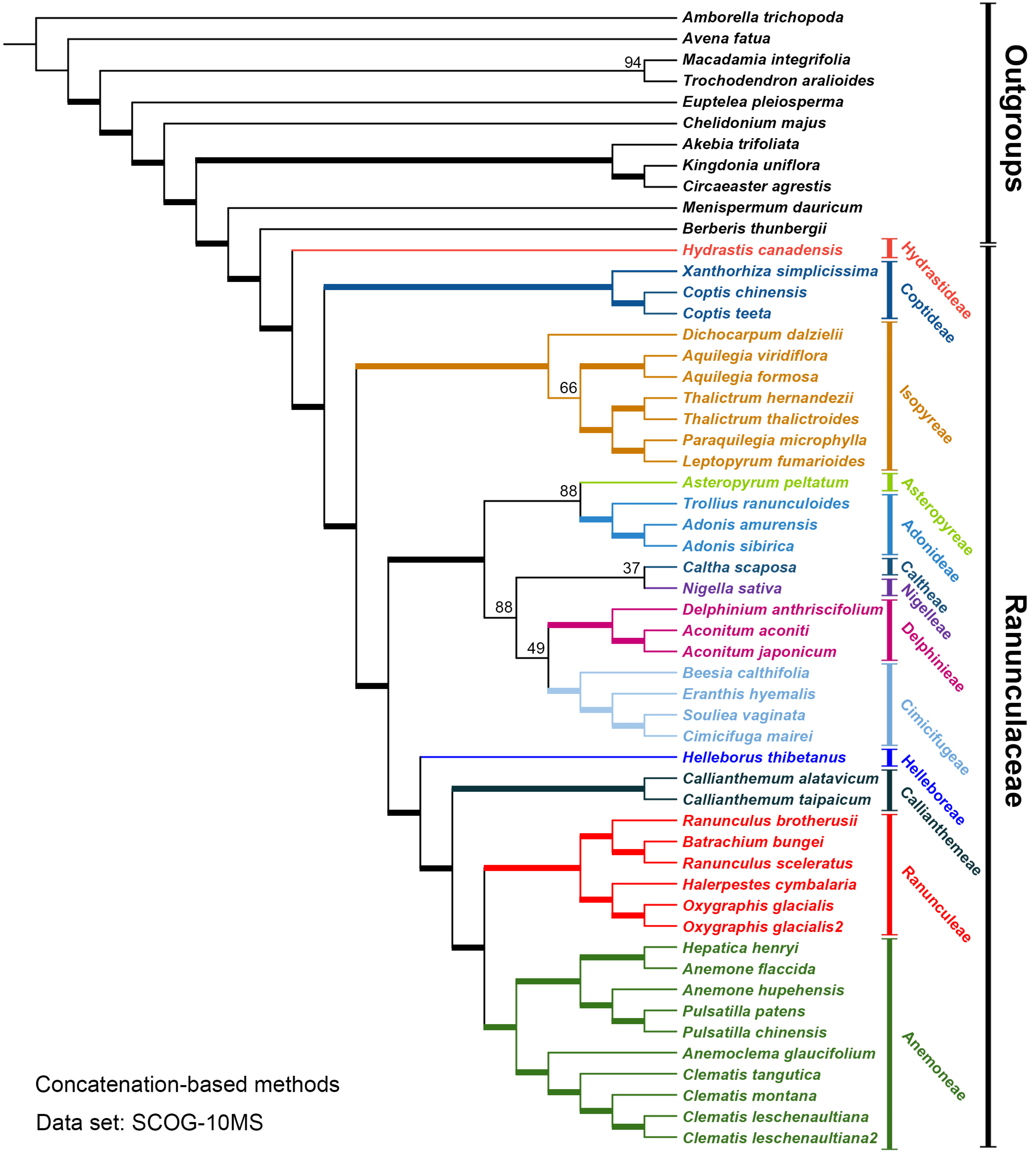

Amborella trichopoda
Avena fatua
94
Macadamia integrifolia
Trochodendron aralioides
Euptelea pleiosperma
Chelidonium majus
Akebia trifoliata
Kingdonia uniflora
Circaeaster agrestis
Menispermum dauricum
Berberis thunbergii
Hydrastis canadensis
Xanthorhiza simplicissima
Coptis chinensis
Coptis teeta
Dichocarpum dalzielii
Aquilegia viridiflora
Aquilegia formosa
66
Thalictrum hernandezii
Thalictrum thalictroides
Paraquilegia microphylla
Leptopyrum fumarioides
Asteropyrum peltatum
88
Trollius ranunculoides
Adonis amurensis
Adonis sibirica
37
Caltha scaposa
Nigella sativa
88
Delphinium anthriscifolium
Aconitum aconiti
49
Aconitum japonicum
Beesia calthifolia
Eranthis hyemalis
Souliea vaginata
Cimicifuga mairei
Helleborus thibetanus
Callianthemum alatavicum
Callianthemum taipaicum
Ranunculus brotherusii
Batrachium bungei
Ranunculus sceleratus
Halerpestes cymbalaria
Oxygraphis glacialis
Oxygraphis glacialis2
Hepatica henryi
Anemone flaccida
Anemone hupehensis
Pulsatilla patens
Pulsatilla chinensis
Anemoclema glaucifolium
Clematis tangutica
Clematis montana
Clematis leschenaultiana
Clematis leschenaultiana2
Hydrastideae
Coptideae
Isopyreae
Asteropyreae
Adonideae
Caltheae
Nigelleae
Delphinieae
Cimicifugeae
Helleboreae
Callianthemeae
Ranunculeae
Anemoneae
Outgroups
Ranunculaceae
Concatenation-based methods
Data set: SCOG-10MS

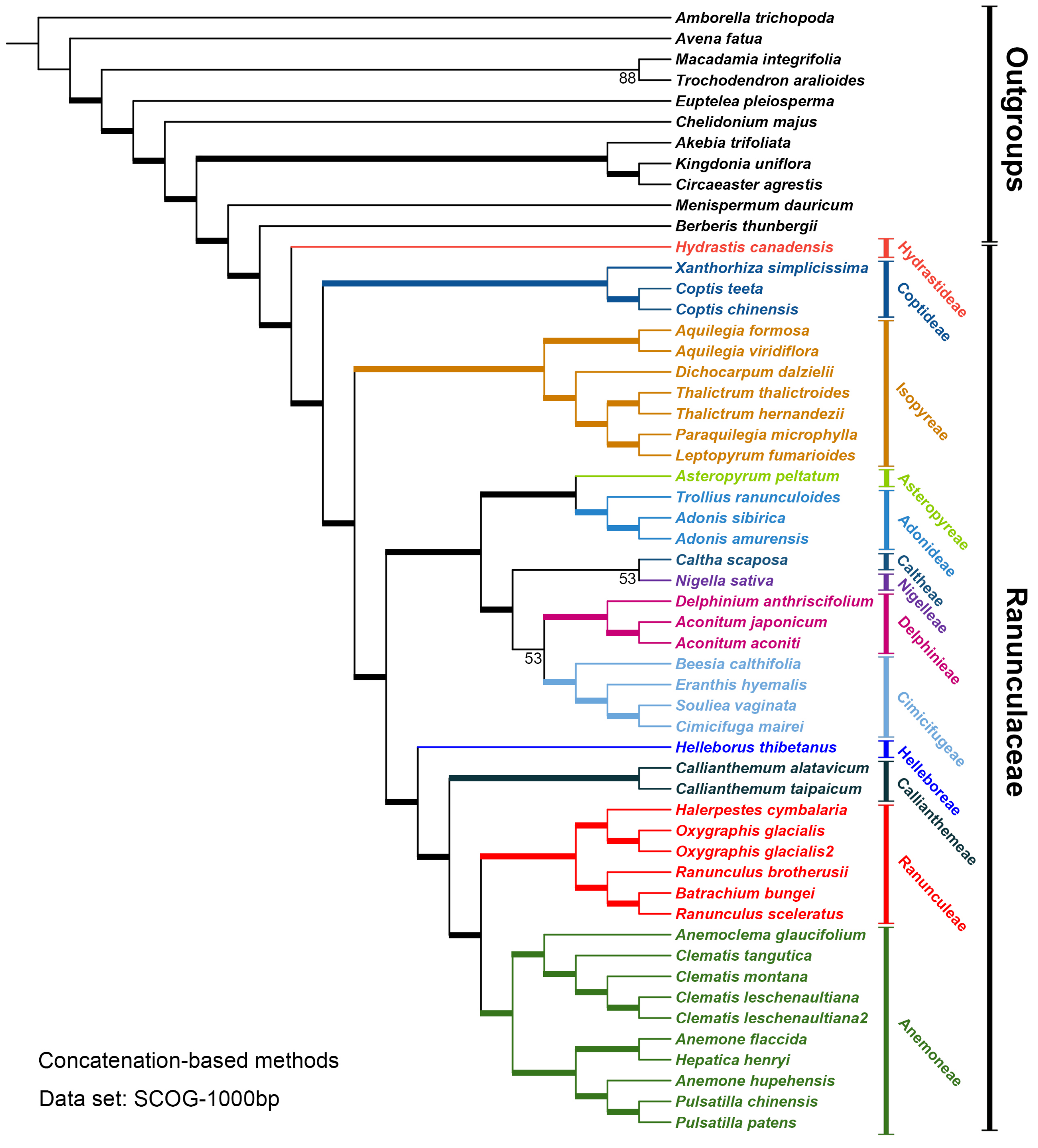

Amborella trichopoda
Avena fatua
Macadamia integrifolia
88
Trochodendron aralioides
Euptelea pleiosperma
Chelidonium majus
Akebia trifoliata
Kingdonia uniflora
Circaeaster agrestis
Menispermum dauricum
Berberis thunbergii
Hydrastis canadensis
Xanthorhiza simplicissima
Coptis teeta
Coptis chinensis
Aquilegia formosa
Aquilegia viridiflora
Dichocarpum dalzielii
Thalictrum thalictroides
Thalictrum hernandezii
Paraquilegia microphylla
Leptopyrum fumarioides
Asteropyrum peltatum
Trollius ranunculoides
Adonis sibirica
Adonis amurensis
Caltha scaposa
53
Nigella sativa
Delphinium anthriscifolium
Aconitum japonicum
Aconitum aconiti
53
Beesia calthifolia
Eranthis hyemalis
Souliea vaginata
Cimicifuga mairei
Helleborus thibetanus
Callianthemum alatavicum
Callianthemum taipaicum
Halerpestes cymbalaria
Oxygraphis glacialis
Oxygraphis glacialis2
Ranunculus brotherusii
Batrachium bungei
Ranunculus sceleratus
Anemoclema glaucifolium
Clematis tangutica
Clematis montana
Clematis leschenaultiana
Clematis leschenaultiana2
Anemone flaccida
Hepatica henryi
Anemone hupehensis
Pulsatilla chinensis
Pulsatilla patens
Outgroups
Ranunculaceae
Hydrastideae
Coptideae
Isopyreae
Asteropyreae
Adonideae
Caltheae
Nigelleae
Delphinieae
Cimicifugeae
Helleboreae
Callianthemeae
Ranunculeae
Anemoneae
Concatenation-based methods
Data set: SCOG-1000bp

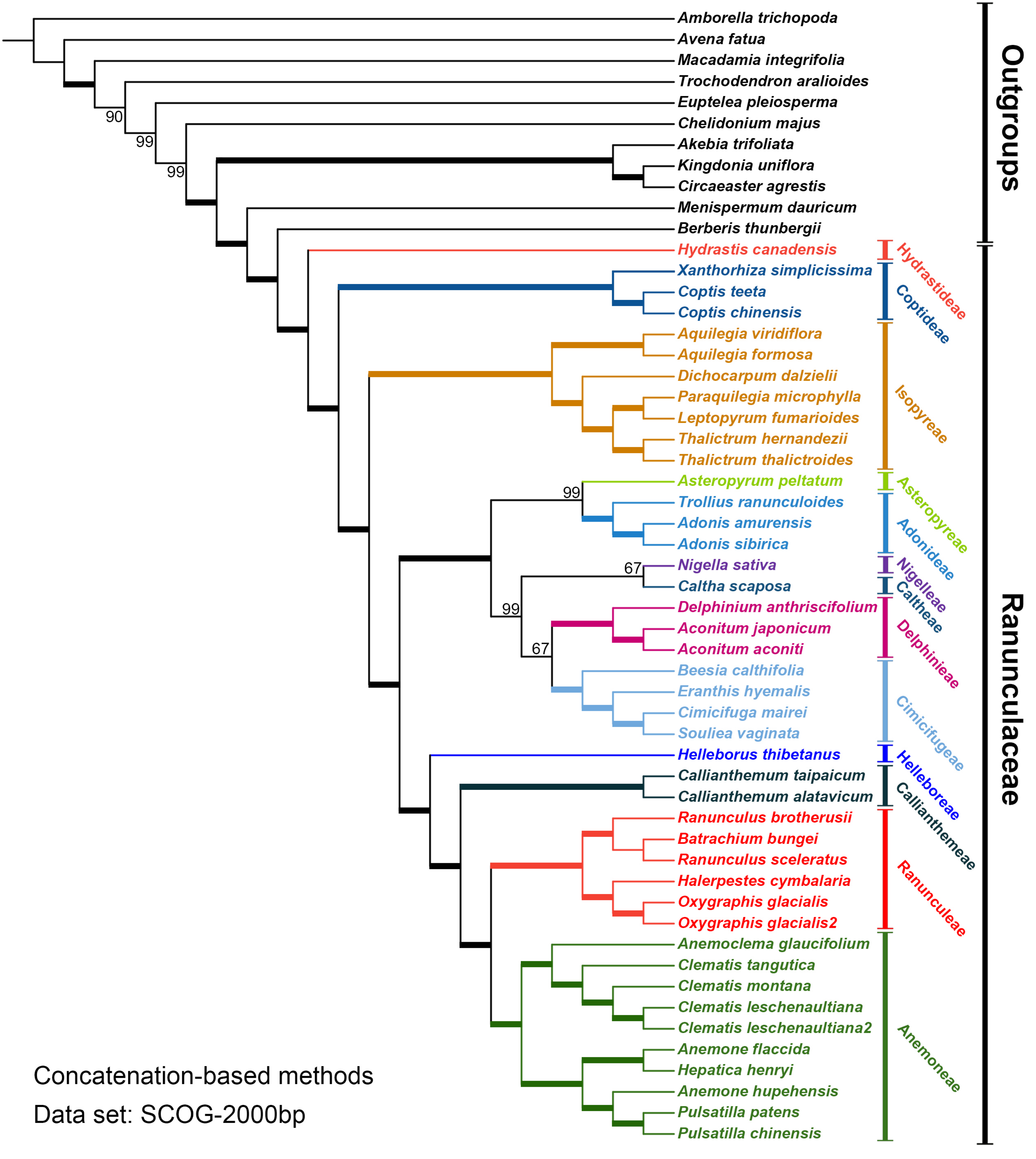

Amborella trichopoda
Avena fatua
Macadamia integrifolia
Trochodendron aralioides
Euptelea pleiosperma
Chelidonium majus
Akebia trifoliata
Kingdonia uniflora
Circaeaster agrestis
Menispermum dauricum
Berberis thunbergii
Hydrastis canadensis
Xanthorhiza simplicissima
Coptis teeta
Coptis chinensis
Aquilegia viridiflora
Aquilegia formosa
Dichocarpum dalzielii
Paraquilegia microphylla
Leptopyrum fumarioides
Thalictrum hernandezii
Thalictrum thalictroides
Asteropyrum peltatum
Trollius ranunculoides
Adonis amurensis
Adonis sibirica
Nigella sativa
Caltha scaposa
Delphinium anthriscifolium
Aconitum japonicum
Aconitum aconiti
Beesia calthifolia
Eranthis hyemalis
Cimicifuga mairei
Souliea vaginata
Helleborus thibetanus
Callianthemum taipaicum
Callianthemum alatavicum
Ranunculus brotherusii
Batrachium bungei
Ranunculus sceleratus
Halerpestes cymbalaria
Oxygraphis glacialis
Oxygraphis glacialis2
Anemoclema glaucifolium
Clematis tangutica
Clematis montana
Clematis leschenaultiana
Clematis leschenaultiana2
Anemone flaccida
Hepatica henryi
Anemone hupehensis
Pulsatilla patens
Pulsatilla chinensis
90
99
99
99
67
99
67
Outgroups
Ranunculaceae
Hydrastideae
Coptideae
Isopyreae
Asteropyreae
Adonideae
Nigelleae
Caltheae
Delphinieae
Cimicifugeae
Helleboreae
Callianthemeae
Ranunculeae
Anemoneae
Concatenation-based methods
Data set: SCOG-2000bp

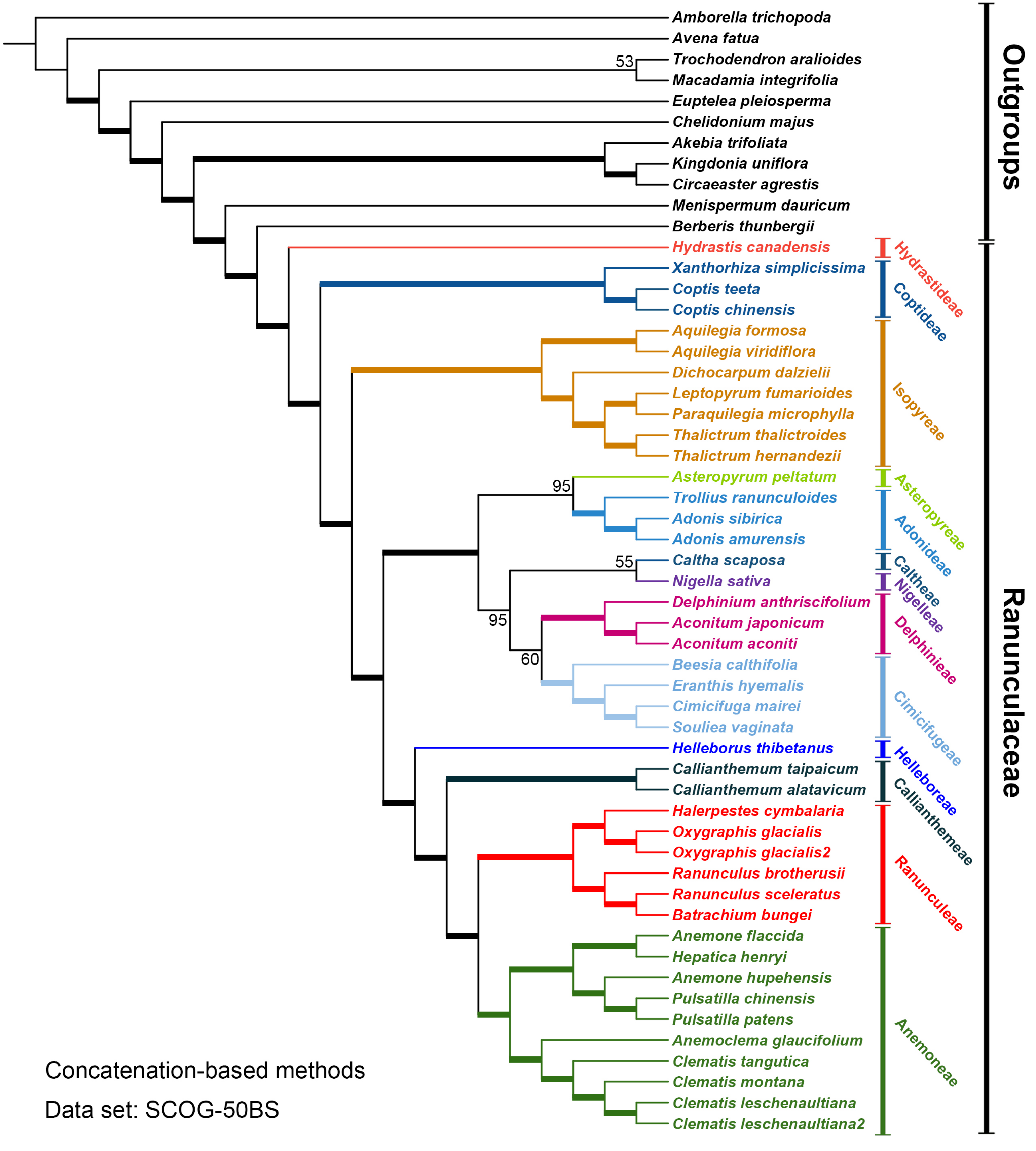
Amborella trichopoda
Avena fatua
53
Trochodendron aralioides
Macadamia integrifolia
Euptelea pleiosperma
Chelidonium majus
Akebia trifoliata
Kingdonia uniflora
Circaeaster agrestis
Menispermum dauricum
Berberis thunbergii
Hydrastis canadensis
Xanthorhiza simplicissima
Coptis teeta
Coptis chinensis
Aquilegia formosa
Aquilegia viridiflora
Dichocarpum dalzielii
Leptopyrum fumarioides
Paraquilegia microphylla
Thalictrum thalictroides
Thalictrum hernandezii
95
Asteropyrum peltatum
Trollius ranunculoides
Adonis sibirica
Adonis amurensis
55
Caltha scaposa
Nigella sativa
95
Delphinium anthriscifolium
Aconitum japonicum
Aconitum aconiti
60
Beesia calthifolia
Eranthis hyemalis
Cimicifuga mairei
Souliea vaginata
Helleborus thibetanus
Callianthemum taipaicum
Callianthemum alatavicum
Halerpestes cymbalaria
Oxygraphis glacialis
Oxygraphis glacialis2
Ranunculus brotherusii
Ranunculus sceleratus
Batrachium bungei
Anemone flaccida
Hepatica henryi
Anemone hupehensis
Pulsatilla chinensis
Pulsatilla patens
Anemoclema glaucifolium
Clematis tangutica
Clematis montana
Clematis leschenaultiana
Clematis leschenaultiana2
Outgroups
Ranunculaceae
Hydrastideae
Coptideae
Isopyreae
Asteropyreae
Adonideae
Caltheae
Nigelleae
Delphinieae
Cimicifugeae
Helleboreae
Callianthemeae
Ranunculeae
Anemoneae
Concatenation-based methods
Data set: SCOG-50BS

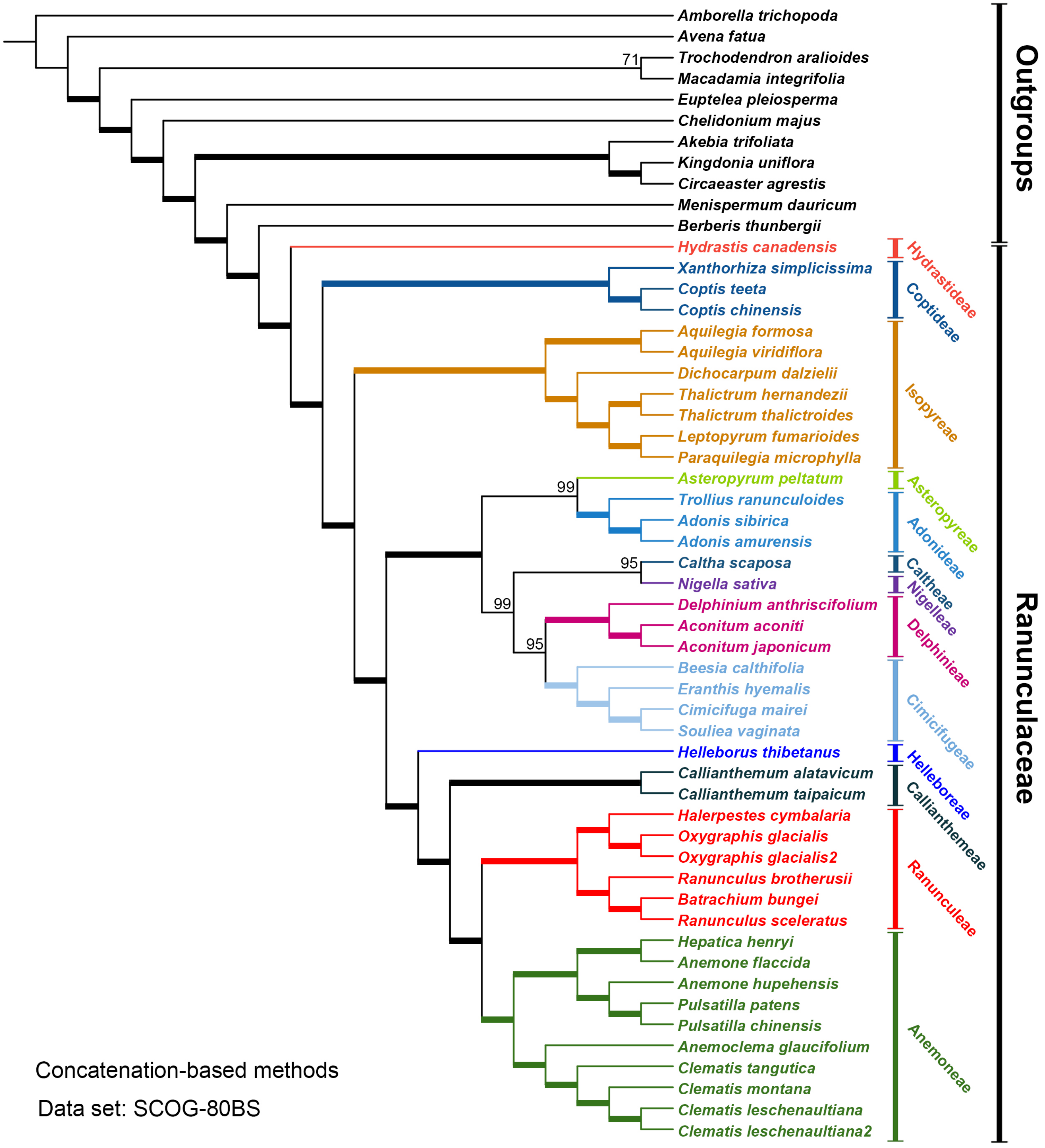

Amborella trichopoda
Avena fatua
71
Trochodendron aralioides
Macadamia integrifolia
Euptelea pleiosperma
Chelidonium majus
Akebia trifoliata
Kingdonia uniflora
Circaeaster agrestis
Menispermum dauricum
Berberis thunbergii
Hydrastis canadensis
Xanthorhiza simplicissima
Coptis teeta
Coptis chinensis
Aquilegia formosa
Aquilegia viridiflora
Dichocarpum dalzielii
Thalictrum hernandezii
Thalictrum thalictroides
Leptopyrum fumarioides
Paraquilegia microphylla
99
Asteropyrum peltatum
Trollius ranunculoides
Adonis sibirica
Adonis amurensis
95
Caltha scaposa
Nigella sativa
99
Delphinium anthriscifolium
Aconitum aconiti
95
Aconitum japonicum
Beesia calthifolia
Eranthis hyemalis
Cimicifuga mairei
Souliea vaginata
Helleborus thibetanus
Callianthemum alatavicum
Callianthemum taipaicum
Halerpestes cymbalaria
Oxygraphis glacialis
Oxygraphis glacialis2
Ranunculus brotherusii
Batrachium bungei
Ranunculus sceleratus
Hepatica henryi
Anemone flaccida
Anemone hupehensis
Pulsatilla patens
Pulsatilla chinensis
Anemoclema glaucifolium
Clematis tangutica
Clematis montana
Clematis leschenaultiana
Clematis leschenaultiana2
Hydrastideae
Coptideae
Isopyreae
Asteropyreae
Adonideae
Caltheae
Nigelleae
Delphinieae
Cimicifugeae
Helleboreae
Callianthemeae
Ranunculeae
Anemoneae
Outgroups
Ranunculaceae
Concatenation-based methods
Data set: SCOG-80BS
