## Supplementary figures and images for "A phylotranscriptome study using silica gel-dried leaf tissues produces an updated robust phylogeny of Ranunculaceae" (from another version of this preprint: version 1, 2021-07-29)

### FIGURE S2

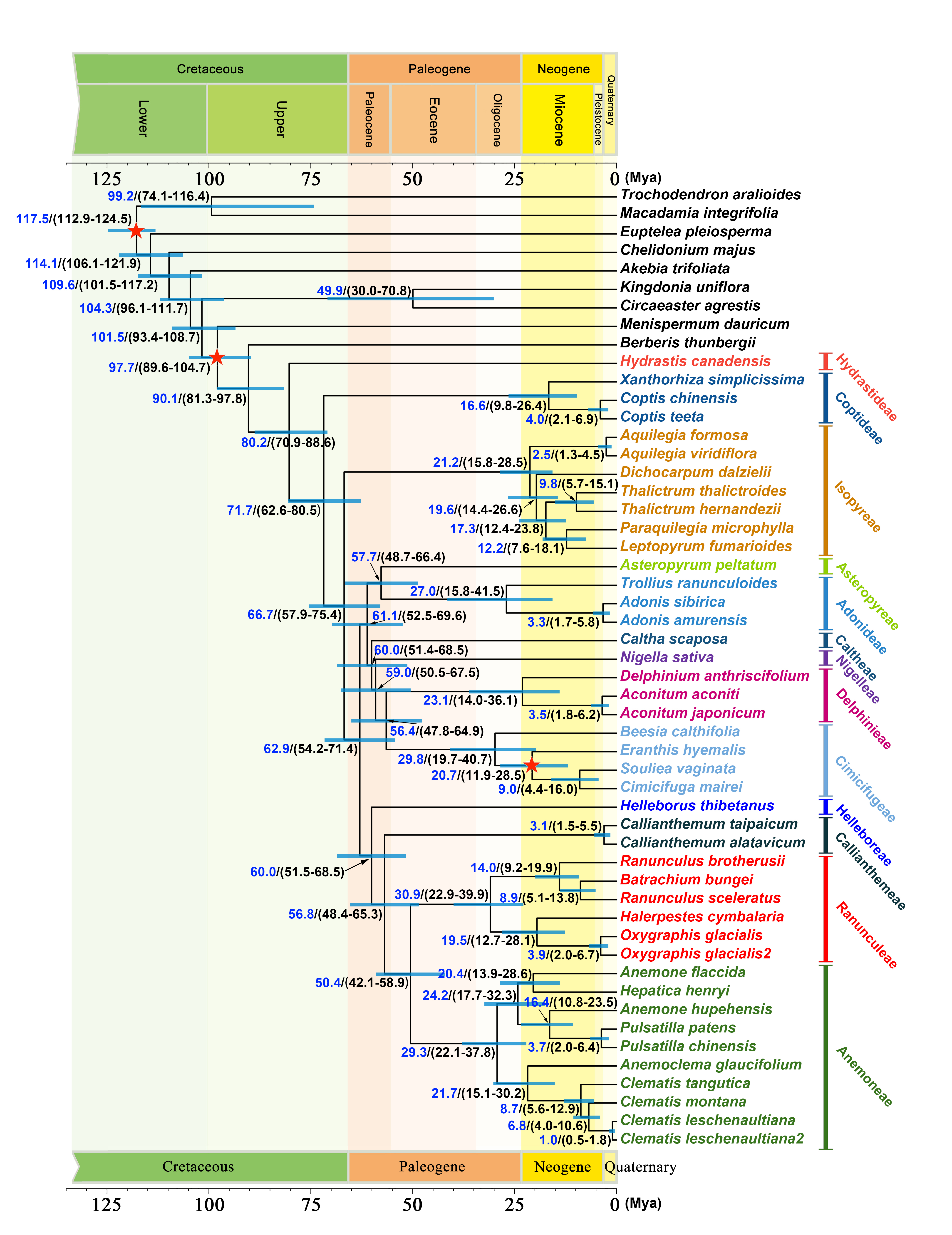
